# The IICR and the non-stationary structured coalescent: demographic inference with arbitrary changes in population structure

**DOI:** 10.1101/341750

**Authors:** Willy Rodríguez, Olivier Mazet, Simona Grusea, Simon Boitard, Lounès Chikhi

## Abstract

In the last years, a wide range of methods allowing to reconstruct past population size changes from genome-wide data have been developed. At the same time, there has been an increasing recognition that population structure can generate genetic data similar to those produced under models of population size change. Recently, Mazet et al. (2016) showed that, for any model of population structure, it is always possible to find a panmictic model with a particular function of population size changes, having exactly the same distribution of *T*_2_ (the coalescence time for a sample of size two) to that of the structured model. They called this function IICR (Inverse Instantaneous Coalescence Rate) and showed that it does not necessarily correspond to population size changes under non panmictic models. Besides, most of the methods used to analyse data under models of population structure tend to arbitrarily fix that structure and to minimise or neglect population size changes. Here we extend the seminal work of Herbots (1994) on the structured coalescent and propose a new framework, the Non-Stationary Structured Coalescent (NSSC) that incorporates demographic events (changes in gene flow and/or deme sizes) to models of nearly any complexity. We show how to compute the IICR under a wide family of stationary and non-stationary models. As an example we address the question of human and Neanderthal evolution and discuss how the NSSC framework allows to interpret genomic data under this new perspective.

**Author summary:** Genomic data are becoming available for a rapidly increasing number of species, and contain information about their recent evolutionary history. If we wish to understand how they expanded, contracted or admixed as a consequence of recent and ancient environmental changes, we need to develop general inferential methods. Currently, demographic inference is either done assuming that a species is a single panmictic population or using arbitrary structured models. We use the concept of IICR (Inverse of the Instantaneous Coalescence Rate) together with Markov chains theory to develop a general inferential framework which we call the Non-Stationary Structured Coalescent and apply it to explain human and Neanderthal genomic data in a single structured model.

## 1 Introduction

Reconstructing the demographic history of populations and species remains one of the great challenges of population genetics and statistical inference (Harpending and Rogers, 2000; Beaumont et al., 2002; Goldstein and Chikhi, 2002; Hey and Machado, 2003; Li and Durbin, 2011; Liu and Fu, 2015; Scerri et al., 2018). In the last decades significant progress has been made in the development of likelihood and likelihood-free methods, hence facilitating the estimation of parameters of interest such as migration or admixture rates, and the dates of putative bottlenecks, expansions or splitting events (Beaumont, 1999; Beaumont et al., 2002; Marjoram et al., 2003; Hey and Nielsen, 2004; Gutenkunst et al., 2009; Li and Durbin, 2011; Bunnefeld et al., 2015).

The rich body of methods and approaches that have been developed during that period can be divided into methods that ignore population structure and thus view the demographic history of species as a series of population size changes (Beaumont, 1999; Chevalet and Nikolic, 2010; Li and Durbin, 2011; Liu and Fu, 2015; Bunnefeld et al., 2015) and those that account for population structure (Nielsen and Wakeley, 2001; Chikhi et al., 2001; Hey and Nielsen, 2004; Gutenkunst et al., 2009; Gronau et al., 2011). In the first family of models, the number of population size changes can be fixed (Beaumont, 1999) or it can itself be estimated (Li and Durbin, 2011; Nikolic and Chevalet, 2014; Liu and Fu, 2015; Boitard et al., 2016). In the second, the model of population structure is typically fixed *a priori* and relatively simple, and its parameters estimated (Chikhi et al., 2001; Hey and Nielsen, 2004; Gutenkunst et al., 2009; Gronau et al., 2011). Some recent methods allow for complex multi-population split models with admixture (Gutenkunst et al., 2009; Gronau et al., 2011). However, while the sizes of ancestral and derived populations can be different in these methods, each one is usually assumed constant and the model structure remains fixed. If the hypothesis made by the underlying models are violated (for example, if the populations evolve under a different type of structure), the estimated parameters may be difficult to interpret (Mazet et al., 2016; Chikhi et al., 2018).

There is thus no general inferential framework that allows the joint estimation of population structure and population size changes (Scerri et al., 2018). This is understandable because it would probably be beyond the current methods to estimate the parameters of such complex models. Still, if we wish to understand the recent evolutionary history of species, including that of humans, it may be necessary to identify the models with or without population structure that can (and those that cannot) explain patterns of genomic diversity.

This is challenging because an increasing number of studies have shown that population structure *per se* can generate spurious signals of population size change in genetic or genomic data. This suggests that the first group of methods may generate misleading histories of population size change (Wakeley, 1999; Storz and Beaumont, 2002; Chikhi et al., 2010; Heller et al., 2013; Mazet et al., 2016; Chikhi et al., 2018) that can explain the data as well as more realistic models of population structure.

Since many species are *de facto* structured in space, a powerful approach to improve the inferential process might be to reduce the model and parameter space so as to focus on models that can explain the data in their genomic complexity. Models that cannot explain the data could then be rejected. For instance, Chikhi et al. (2018) showed, using simulated IICR (inverse instantaneous coalescence rate) plots defined in Mazet et al. (2016), that several models used to quantify admixture between humans and Neanderthals cannot explain human and Neanderthal PSMC plots (Li and Durbin, 2011).

In the present study we introduce a mathematical and conceptual framework based on the structured coalescent (Herbots, 1994). We show that IICR curves can be used to develop a powerful model choice and model exclusion strategy for structured models of nearly any complexity. In a few words, the IICR is a time-dependent function that can be interpreted as an effective size in a panmictic population. However, for structured models this interpretation may be misleading. For instance, there are various IICR curves for the same demographic model that depend on the temporal and geographical sampling scheme (Mazet et al., 2016; Chikhi et al., 2018). IICR curves can thus be seen as sample-dependent coalescent histories, which together may represent a unique signature for a complex model. The IICR is related to the PSMC method of Li and Durbin (2011) in the sense that the PSMC method, while generally interpreted in terms of population size changes, actually infers the IICR for a sample of size two (Mazet et al., 2016). The IICR curves can thus be seen as summaries of genomic information (Chikhi et al., 2018).

We extend previous work on the IICR by applying the theory of Markov chains (see for example Norris (1998)) to models of population structure of nearly any complexity (i.e., including changes in gene flow and/or deme sizes). We show how the transition rate matrices associated to a given structured model can be used to compute the corresponding IICR curves with very high accuracy, with a much lower computational time than the simulation-based approach used in Chikhi et al. (2018). We apply this new framework to the structured coalescent of Herbots (1994) and extend it to non-stationary models, hence introducing the Non-Stationary Structured Coalescent (NSSC), and discuss the possibility to extend it to less constrained genealogical models.

To that aim we first review and summarise the main results and terminology required to link the Markov chain described by the structured coalescent with the notion of IICR. We acknowledge the seminal work of Herbots (1994) who derived the transition rate matrix corresponding to the structured coalescent. We apply this approach to compute the IICR of several models of population structure, such as the *n*-island model, and 1D and 2D stepping stone models, under arbitrary sampling schemes. Using the semi-group property we show how our results can be naturally extended to models with an arbitrary number of changes in gene flow. We then show how demes with different sizes (e.g., continent-island models), or changes in the deme sizes can be easily incorporated into this framework. In addition, we show that transition rate matrices can be simplified using symmetries for several models (n-island, continent-island) reducing the computational costs by several orders of magnitude. We finally apply these results to humans and Neanderthals and identify models of population structure that can explain human and Neanderthal genomic diversity.

### 2 The structured coalescent and transition rate matrices: towards the IICR

The distribution of coalescence times in models that account for population structure (i.e., population subdivision) has been the centre of interest of important and early theoretical studies (Takahata, 1988; Notohara, 1990; Herbots, 1994; Barton and Wilson, 1995; Wakeley, 1999, 2001; Nordborg, 2001; Charlesworth et al., 2003). In particular, Herbots (1994) developed an elegant extension of the coalescent (Kingman, 1982) for structured populations under a number of constraints regarding gene flow (see below). This extension, named structured coalescent, has been extremely important and is based on a continuous-time Markov chain. It allows to compute explicitly the moment-generating function of the coalescence times under a wide range of models considering population structure (Herbots, 1994; Wilkinson-Herbots, 1998). In this section we review the terminology and theory leading to the structured coalescent, introduce transition rate matrices and show how they can be used to compute the IICR of Mazet et al. (2016).

### 2.1 From the discrete-time model to the continuous-time approximation

Following Herbots (1994), we consider a haploid population divided into a finite number *n* of subpopulations or demes which are panmictic and whose size, *N_i_* for deme *i,* is assumed to be large. Each deme is also assumed to behave as a haploid Wright-Fisher model. These demes are connected to each other by migration events. Every generation a proportion *q_ij_* of the haploid individuals from deme *i* migrates to deme *j* (migrants are chosen without replacement, independently and uniformly from deme *i*). We assume that deme sizes and migration rates are constant in time. In this model the number of haploid individuals in deme *i* is *N_i_* = 2*c_i_N,* where *c_i_* is a positive integer and *N* is large. Also, the proportion *q_ij_* is of the order of *1/N* for every (*i*, *j*). In the classical n-island model of Wright (1931), the *c_i_* are all identical and set to one. If we set 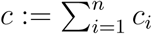, we can write the total haploid population size as *N_T_* = *2cN.* Note that in diploid applications, *c_i_N* is the number of diploid individuals in deme *i* and thus the diploid population size will be *cN.*

The structured coalescent of Herbots (1994) assumes that the size of each subpopulation is maintained constant under migration, which generates the following constraint at the population level:

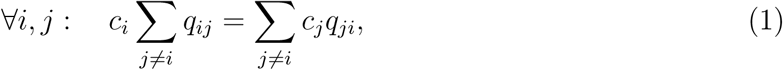

where *q_ij_* is the probability that one individual migrates from deme *i* to deme *j.* In other words, all outward migrants must be replaced by inward migrants from the other islands.

This condition is required in the structured coalescent but we stress that it is not required in the structured model of Notohara (1990) or when simulating data under structured models using the *ms* software of Hudson (2002).

Looking now backward in time, Herbots (1994) defines the *backward migration parameter* from deme *i* to deme *j* (denoted *m_ij_*) as:

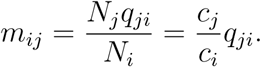

The backward migration parameter *m_ij_* represents the proportion of individuals in deme *i* that were in deme *j* just before the migration step. Also, 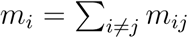 represents the proportion of individuals inside deme *i* that were in a different deme just before the migration step.

In this backward perspective, we suppose that we have a sample of *k* haploid genomes at a time which we arbitrarily call time zero. We then trace back the ancestral history of the *k* lineages until their MRCA (Most Recent Common Ancestor). We are interested in the statistical properties of the gene trees of this sample of *k* lineages at different loci in the genome. Following Herbots (1994), we define *α_N_* := {*α_N_*(*r*); *r* = 0, 1, 2,…}, where *α_N_*(*r*) is a vector whose *i*^th^ component denotes the number of distinct lineages in subpopulation *i, r* generations ago.

Herbots (1994) proved that, measuring time in units of 2*N* generations, *α_N_* converges to a continuous-time Markov chain called the structured coalescent, as *N* tends to infinity and as all *m_ij_* (*i* ≠ *j*) tend to zero, in such a way that *M_ij_*/2 := 2*Nm_ij_* and 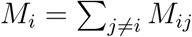 are constant, finite and non-zero. In the rest of the manuscript we drop the *N* index in *α_N_,* but we wish to stress that *α_N_*(*r*) represents the configuration of the remaining ancestral lineages at generation *r* backwards in the discrete-time model and *α*(*t*) represents the ancestral configuration *t* time units ago, in the continuous-time model. When *r* = 0 or *t* = 0, it is simply the initial sample configuration. The structured coalescent is thus the continuous-time Markov chain whose states are all the possible configurations for the ancestral lineages at different times in the past. It is thus characterised by the transition probabilities between configurations. A key element describing this Markovian process is its transition rate matrix denoted *Q* hereafter.

### 2.2 The transition rate matrix of a continuous-time Markov chain

Transition rate matrices are briefly introduced in this section. For a full background, see for instance Norris (1998).

A transition rate matrix on the finite set *I* is a square matrix *Q* = *Q*(*i, j*), with *i, j* ∊ *I* satisfying the two following conditions:

- ∀*i* ≠*j*, *Q*(*i*, *j*) ≥ 0,
- 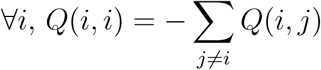.

If we now define, for all *t* ≥ 0, the exponential matrix *P_t_* = *e^tQ^* which has the same size as *Q*, *P_t_* then satisfies the following properties, for all *s, t:*

- *P_t+s_* = *P_t_ P_s_* (semigroup property),
- 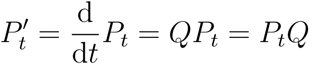,
- *P_t_* is a stochastic matrix (each coefficient is non-negative and the sum over each row is one).

Also each coefficient of the matrix *P_t_,* for all *t* ≥ 0, is a transition probability:

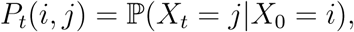

where (*X_t_*)*_t_*_≥0_ is a continuous-time Markov chain on the finite set *I.* In other words, *X_t_* is a jump process, whose behaviour is the following:

- if at a given time *s* ≥ 0 we have *X_s_* = *i,* then it jumps away from state *i* after an exponential time of parameter −*Q*(*i*, *i*), which does not depend on *s,*
- at each jump from state *i,* the rate at which state *j* is reached is 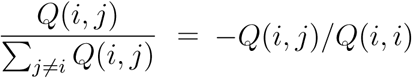.

The transition rate matrix *Q* then contains all the information on the behaviour of (*X_t_*)*_t_*_≥0_, given the initial condition *X*_0_. We can see that, for all *i* ∊ *I,* the parameter *Q*(*i*, *j*) is *the rate of going from i to j,* as soon as *j* ≠ *i,* and the parameter −*Q*(*i*, *i*) is the *rate of leaving i.*

In the case of the *structured coalescent* the jump process of interest is the *ancestral lineage process.* The set *I* is the set of possible configurations *α* = (*α*_1_, …, *α_n_*), where *α_i_* is the number of lineages present in the *i*^th^ deme, and *n* the number of demes. A “jump” between two configurations occurs when a lineage migrates from one deme to another (say, from deme *i* to deme *j*), or when a coalescence takes place within a deme in which there are at least two lineages. We thus have now all the elements necessary to compute the IICR for stationary models under the structured coalescent.

## 3 Transition rate matrices allow us to compute the IICR for a wide family of structured models

Mazet et al. (2016) introduced and defined the IICR. They derived it analytically for the *n-*island model and for *k* = 2 lineages for the only two distinguishable sampling schemes (the two lineages in the same deme, respectively in different demes) available for that model (initial configurations or states of the Markov chain). In this section we show how transition rate matrices can be used to analyse a wide family of models of population structure. We take the case of *k* = 2 and step by step explain how the transition rate matrix can be constructed. We then describe the general algorithm used to construct the IICR for all the models analysed here for *k* = 2. We finally apply this method to the n-island model and show that we can re-derive the results obtained by Mazet et al. (2015) and Herbots (1994).

### 3.1 General case

As noted above, Herbots’ discrete-time process converges to a continuous-time Markov process (called structured coalescent). Here we describe in more details how to construct the associated transition rate matrix. Let’s assume that we have numbered the demes of the model from 1 to *n* where *n* is the total number of demes. Then the vector (*c*_1_, *c*_2_, *…*, *c_n_*) indicates the size of each deme. We take a sample of *k* genes (*k* ≥ 2) from the population at the present (*t* = 0) and we trace the ancestral lineages back to the MRCA. The vector *α* = (*α_i_*)_1≤*i*≤*n*_, where *α_i_* is the number of ancestral lineages in deme *i,* represents a possible ancestral configuration for the lineages when going backwards in time. For example, for *n* = 3 demes and *k* = 2 samples, the vector *α* = (1, 1, 0) is an element of 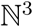 and indicates that there is one ancestral lineage in deme 1, one ancestral lineage in deme 2 and no ancestral lineage in deme 3. Note that 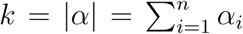. We call *E_k,n_* the set of all possible states of a structured model with *n* demes and a sample of size *k.* We have:

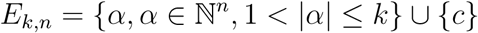

where *c* represents the state when the MRCA of the sample is reached (|*α|* = 1) .

The Markov chain can change from one state *α* ∊ *E_k,n_* to another state *β* ∊ *E_k,n_* either by a migration event (which implies that *|β|* = |*α*|) or by a coalescence event inside a deme (which implies that *|β|* = |*α*| − 1). Before constructing the associated transition rate matrix we need to define an order on *E_k,n._* We choose the inverse lexicographical order. For example for *n* = 3 and *k* = 2 it would be:

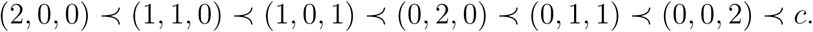

Note that the state *c* (when the MRCA of the sample is reached) is placed in the last position. We denote *ϕ* the function that associates an element of *E_k,n_* with the corresponding index in the inverse lexicographical order. For example, taking the previous example, for *α* = (2, 0, 0) it will be *ϕ*(*α*) = 1 and *ϕ*(*c*) = 7. Throughout the next sections we will assume that there is an order on the set *E_k,n_* given by the function *ϕ.* We define *n_α_* := *ϕ*(*α*) so that *P_t_*(*n_α_*, 1) refers to the first element of the row *n_α_* in the matrix *P_t_*.

The corresponding transition rate matrix can be constructed as:

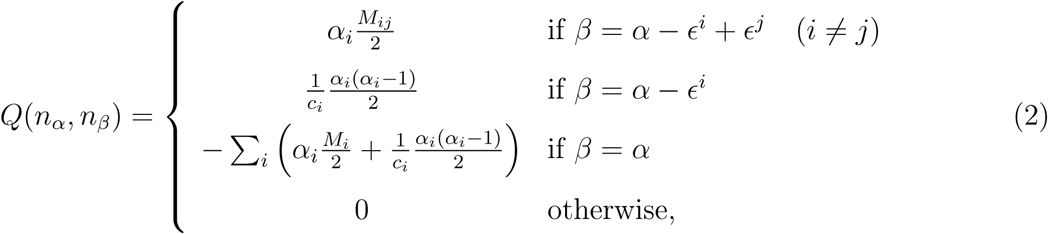

where *ϵ^i^* is the vector whose components are 1 on the *i*^th^ position and 0 elsewhere. The matrix *Q* describes two types of possible events for each configuration *α:*

- *β* = *α* − *ϵ^i^* + *ϵ^j^* when one lineage migrates (backward in time) from island *i* to island *j.* The rate of this migration is *M_ij_*/2 (migration rate to deme *j* for each lineage in deme *i*) times *α_i_,* the number of lineages present in deme *i.*
- *β* = *α* − *ϵ^i^* denotes a coalescence event between two lineages in deme *i,* which reduces the number of lineages by one in this deme. This occurs only if *α_i_* ≥ 2. If this is not the case we can see that *α_i_*(*α_i_* − 1) = 0. The term *α_i_*(*α_i_* − 1)/2 is the number of possible pairs among the *α_i_* lineages. This term is multiplied by 1/*c_i_* since the *i*^th^ island has a population size equal to 2*c_i_N,* and 1*/c_i_* is the coalescence rate for each pair of lineages in this island since time is scaled by 2*N.*

Since no other kind of event can occur than a migration or a coalescence, and multiple coalescences or migrations are negligible, the other rates are null. Note that the opposite of the diagonal coefficient −*Q*(*n_α_, n_α_*) is the total jump rate from configuration *α.*

The transition rate matrix can be very large depending on the model of population structure assumed and on the sample size. For *k* ≤ *n* the number of states is on the order of *n^k^,* and the matrix will have on the order of *n*^2*k*^ terms.

### 3.2 Case of a sample of two lineages (*k* = 2)

We now consider the case where we take a sample of two lineages (i.e., *k* = 2 corresponding to two haploid genomes or one diploid genome) in an arbitrary model of population structure with *n* demes of size 2*c_i_N*, for large *N.* We can reduce all possible configurations to only two types of configurations, excluding the configuration where the two lineages have coalesced:

- both lineages are in the same deme *i: α* = 2*ϵ^i^*,
- the two lineages are in different demes, say, demes *i* and *j* with *i* ≠ *j: α* = *ϵ^i^* + *ϵ^j^*.

When the two lineages are in the same deme (first case), there are two possible events that can change the configuration: a coalescence with rate 1*/c_i_,* or a (backward) migration from *i* to *j* ≠ *i* for each lineage, with rate *α_i_M_ij_*/2 for both lineages, hence a total rate of *α_i_M_ij._* When a coalescence happens, the number of lineages decreases by one. When a migration from deme *i* to deme *j* happens, the new configuration is one in which the lineages are now in different demes, which is a second-type configuration.

When the two lineages are in different demes, no coalescence can occur and the two lineages may either stay in the same deme or migrate to another deme, from *i* to *ℓ* (which can be equal to *i*) for the first lineage, with rate *α_j_M_iℓ_*/2 or from *j* to *ℓ* (which can be equal to *i*) for the second lineage, with rate *α_j_M_jℓ_*/2. If the lineages end up in the same deme we are back to a configuration of the first type, otherwise, we end up in a second-type configuration.

By definition, the number of rows and columns of the full transition rate matrix (that we will call *n_c_*) is the number of different configurations for the ancestral lineages. In the case of a model with *n* demes and a sample of size *k* = 2, we have that *n_c_* = *n*^2^ + 1. We will assume that the “last configuration” is the one in which the two lineages have coalesced, and thus ignore where the coalescence took place. Also note that the rate of a coalescence event in deme *i* (which is equal to 1/*c_i_*) depends on the size of deme *i.* In the transition rate matrices that we will use here the coalescence configuration corresponds to the last row and column.

### 3.3 General algorithm for the construction of the transition rate matrix for *k* = 2

Here we give a general algorithm that can be used to construct the transition rate matrix of a given model. The first step is to explicitly order all the demes. Then, given the number *n* of (ordered) demes the set of all possible configuration for *k* = 2 lineages is:

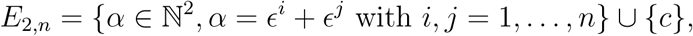

where *ϵ^i^* + *ϵ^j^* means that there is one lineage in deme *i* and one lineage in deme *j* (note that it could be *i* = *j*); and *c* is the configuration where both lineages have coalesced.

As in section 3.1 we take the inverse lexicographical order on *E*_2*,n.*_ Define *ϕ* as a function from *E*_2*,n*_ to {1, 2, …, |*E*_2,*n*_|} such that *ϕ*(*α*) is the index of *α* according to the inverse lexicographical order. Then *ϕ*^−1^ is the inverse of *ϕ* and *ϕ*^−1^(*i*) gives the element of *E*_2*,n*_ which is at position *i* according the inverse lexicographical order.

Once the function *ϕ* is defined and we have the values of *C* = (*c*_1_, … *, c_n_*) (the size of the demes) and *M_ij_* (the migration matrix), we can use the following algorithm to construct the transition rate matrix *Q:*

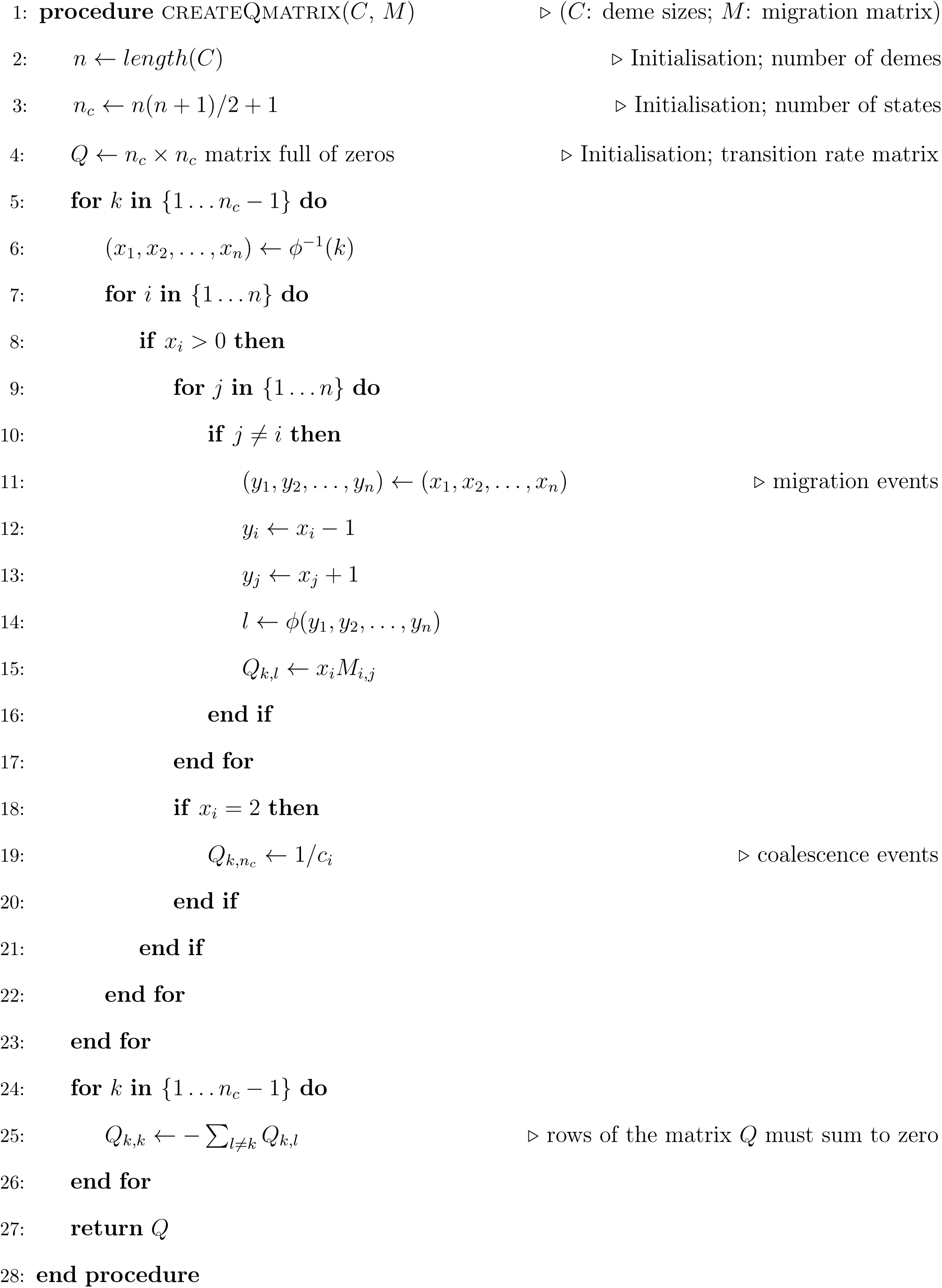

Note that since the last configuration (coalescence) is an absorbing state of the Markov process, the last row has only zeros.

### 3.4 Using the transition rate matrix to derive the distribution of coalescence times and evaluate the IICR for samples of size two

We now focus on the coalescence time between **two** lineages and see that we can derive the IICR in terms of transition rate matrices. The theory of Markov chains (Norris, 1998) gives the tools allowing to compute the probability distribution of *T*_2_ based on the matrix exponential of the transition rate matrix for the model of interest

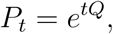

where *P_t_* is the transition semigroup of the corresponding Markov process, i.e., 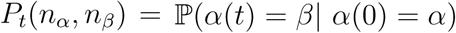, where we *α*(*t*) denotes the ancestral lineages configuration at time *t* in the past and *α*(0) represents the initial sample configuration.

As noted in Section 2.2, the terms of *P_t_* represent the transitions probabilities of interest. For instance, the term in row *n_α_* and column *n_β_* of *P_t_* represents the probability that the process is in the configuration *β* at time *t* given that it was in the configuration *α* at time zero. Thus, the probability that two lineages in the configuration *α* at *t* = 0 have reached their most recent common ancestor at time *t* can be found as *P_t_*(*n_α_, n_c_*), where *n_c_* is the last column since *n_c_* = *ϕ*(*c*) is the column number of the coalescence state.

Consequently, if we denote by 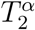 the coalescence time of two lineages sampled in the configuration *α,* the cumulative distribution function (*cdf*) of this random variable can be computed from the transition semigroup:

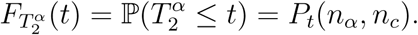

The probability density function (*pdf*) of 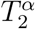, 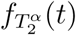, is by definition the derivative of 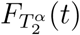. It can thus be computed from the matrix *P_t_,* by using the property

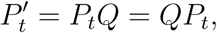

where 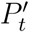 is the matrix whose cells contain the derivative of the corresponding cells of *P_t_.* We can thus write

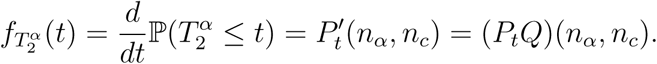

It is then easy to derive, for any time *t* ≥ 0, the instantaneous coalescence rate, which is the probability to coalesce at time *t* given that the lineages have not coalesced yet. This is by definition the ratio

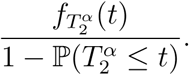

The Inverse Instantaneous Coalescence Rate (IICR) of Mazet et al. (2016), is simply the inverse of this ratio, in which all the terms can be written as a function of *P_t_* and the transition rate matrix, namely:

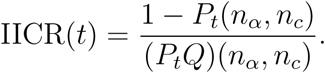

In the next section, we show how transition rate matrices can be used to re-derive the analytical results of Mazet et al. (2016) on the IICR of the *n*-island model.

### 3.5 The IICR of the *n*-island model for *k* = 2 using the simplified transition rate matrices

In the symmetric island model of Wright (1931) the *n* demes (*n* ≥ 2) are equal-sized islands with the same migration rate between any two islands (Figure 2. With the notations above, we have ∀*i* = 1,…,*n, c_i_ =* 1, *M_i_* = *M* and *M_ij_* = *M*/(*n* − 1) for *j* ≠ *i.* Taking into account the fact that the model is fully symmetrical, we only need to consider two configurations for a sample of two lineages: they are either in the *same* deme (denoted *s*) or in *different* demes (denoted *d*). There is a third state that corresponds to the coalescence event which takes place at rate 1. We thus obtain the simplified transition rate matrix

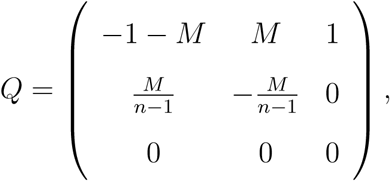

where the first configuration is *s,* the second is *d,* and the third one corresponds to a coalescence event, which can only occur when both lineages are in the same island.

This matrix is simple and small enough to allow the derivation of explicit formula for its exponential *P_t_* = *e^tQ^* and hence for the corresponding IICR functions under the two possible starting configurations (IICR_s_ or IICR_d_ for samples taken in the same or different demes respectively):

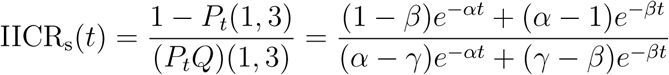

and

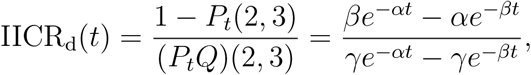

with

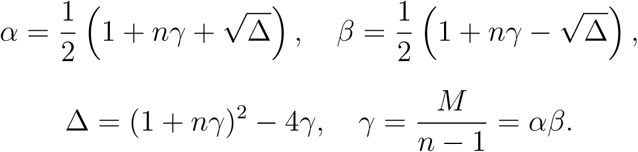

These formulae are identical to those of Mazet et al. (2015), who obtained them using a different approach. We can see the plots of the IICR_s_ and IICR_d_ for the n-island model in Figure 1.

**Figure 1:**
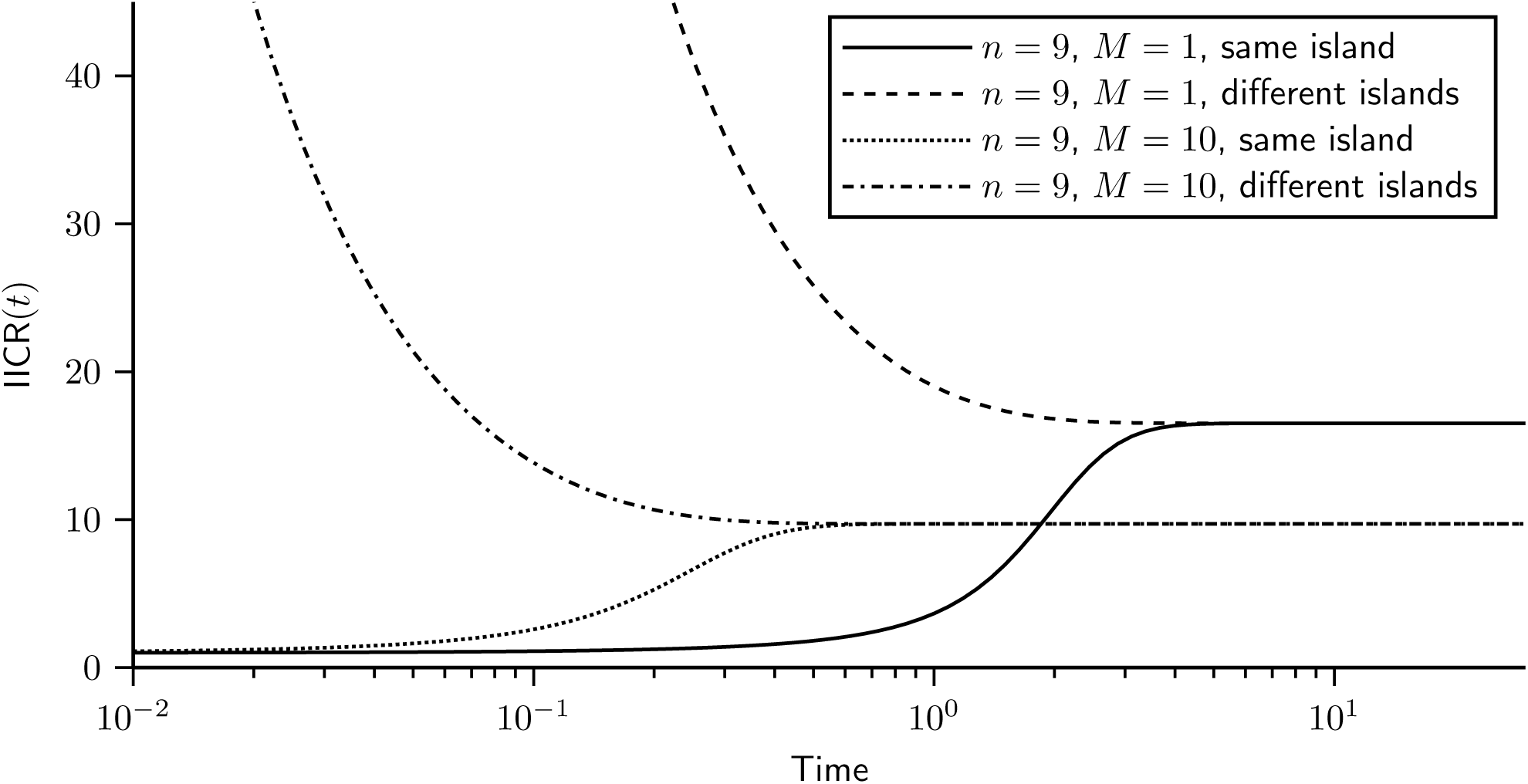
IICR for the n-island model. We plotted the IICR for a model with *n* = 9 islands and assuming two values for the migration rate, *M* = 1 and *M* = 10. For each model we started with the two configurations in which the genes are either sampled in the *same* (IICR_s_) or in *different* (IICR_d_) islands.

**Figure 2:**
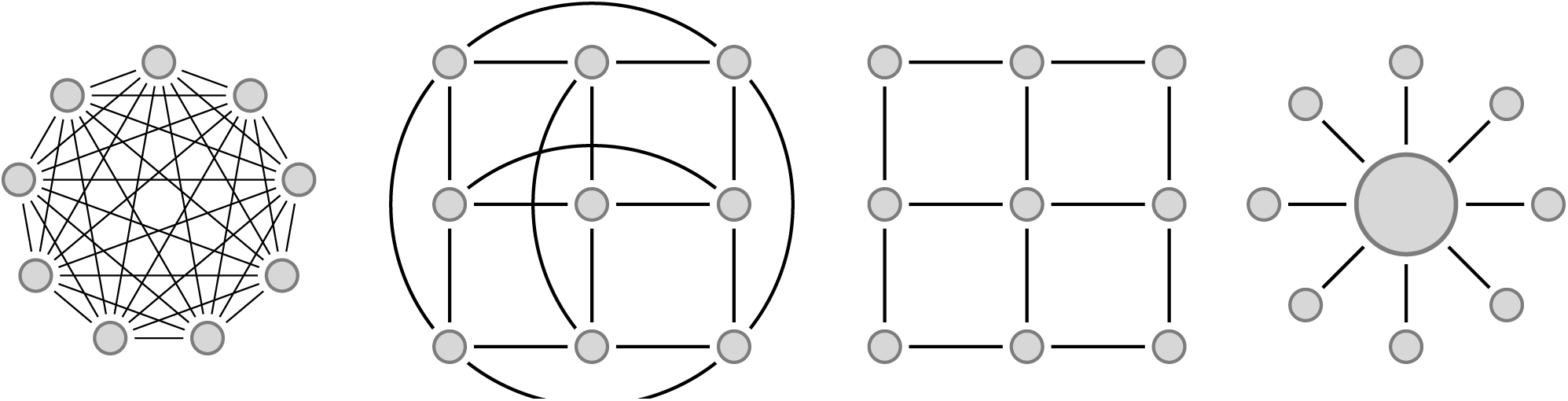
Diagrams for commonly used structured models. From left to right: *n*-islands, torus 2D stepping stone, 2D stepping stone and continent-island model.

## 4 Constructing the IICR for two stationary models, the 2D stepping stone and continent-island models

We now apply the framework and algorithm described above to two stationary models. To our knowledge, there is no analytical expression for the distribution of the coalescence time *T*_2_ under these two models. The transition rate matrices and IICR results for several other stationary models are shown in the Supplementary Materials.

### 4.1 2D stepping stone models with and without edges

Stepping stone models (Kimura, 1994; Malécot and Blaringhem, 1948) assume that the demes are located at the nodes of a regular lattice in one or two dimensions (hereafter 1D and 2D stepping stone models). Each deme can have up to four neighbours and migration events are only possible between neighbouring demes. These models incorporate space, and are thus thought to be more realistic than the *n*-island model described above, which implicitly assumes that migration is as likely between neighbouring than between distant islands. The border demes can either be connected with each other, hence forming a torus, or can behave as bouncing borders (Figure 2). In some models the bouncing borders migrants are assumed to stay in their deme, whereas in other models they are distributed among the demes to which their deme is connected.

For the 2D stepping stone model, we set, ∀*i*, *j* = 1,…, *n*, *c_i_* = 1 and *M_ij_* = *M/*4 if islands *i* and *j* are neighbours, and *M_ij_* = 0 otherwise. The difference between the models with and without edges used here is thus in the way neighbours are defined. In the model with borders the four corner islands have only two neighbours, the islands on the borders of the lattice have three, and the others have four neighbours (see Figure 2).

Figure 3 shows the IICR_s_ (two haploid genomes sampled in the same deme, or one diploid genome), for a 3 × 3 stepping stone model with and without borders (Figure 2). In the latter case (no borders), all demes are statistically identical, and there can thus be only one IICR_S_ plot. In the model with borders, there are three possible ways to sample a diploid individual, and three IICR_s_ are plotted. This figure confirms the results of Chikhi et al. (2018) by showing that the IICR_s_ plots for a stepping stone are also S-shaped. They all start in the recent past at a value equal to the deme size and converge in the ancient past towards the same plateau. However, it is remarkable that they differ in the trajectory from the present to the plateau value, depending on the location of the deme (corner, border or centre). These results thus confirm that in a stepping stone model, two diploid individuals sampled in different demes (i.e., geographical regions) will both exhibit signals of population decrease that will be different even though the population size was constant and they both belonged to the same structured model (Chikhi et al., 2018). Note that, as for the *n*-island model, the IICR exhibits a signal of spurious population increase when the two genes are sampled in different demes (IICR_d_, see Supplementary Materials).

**Figure 3:**
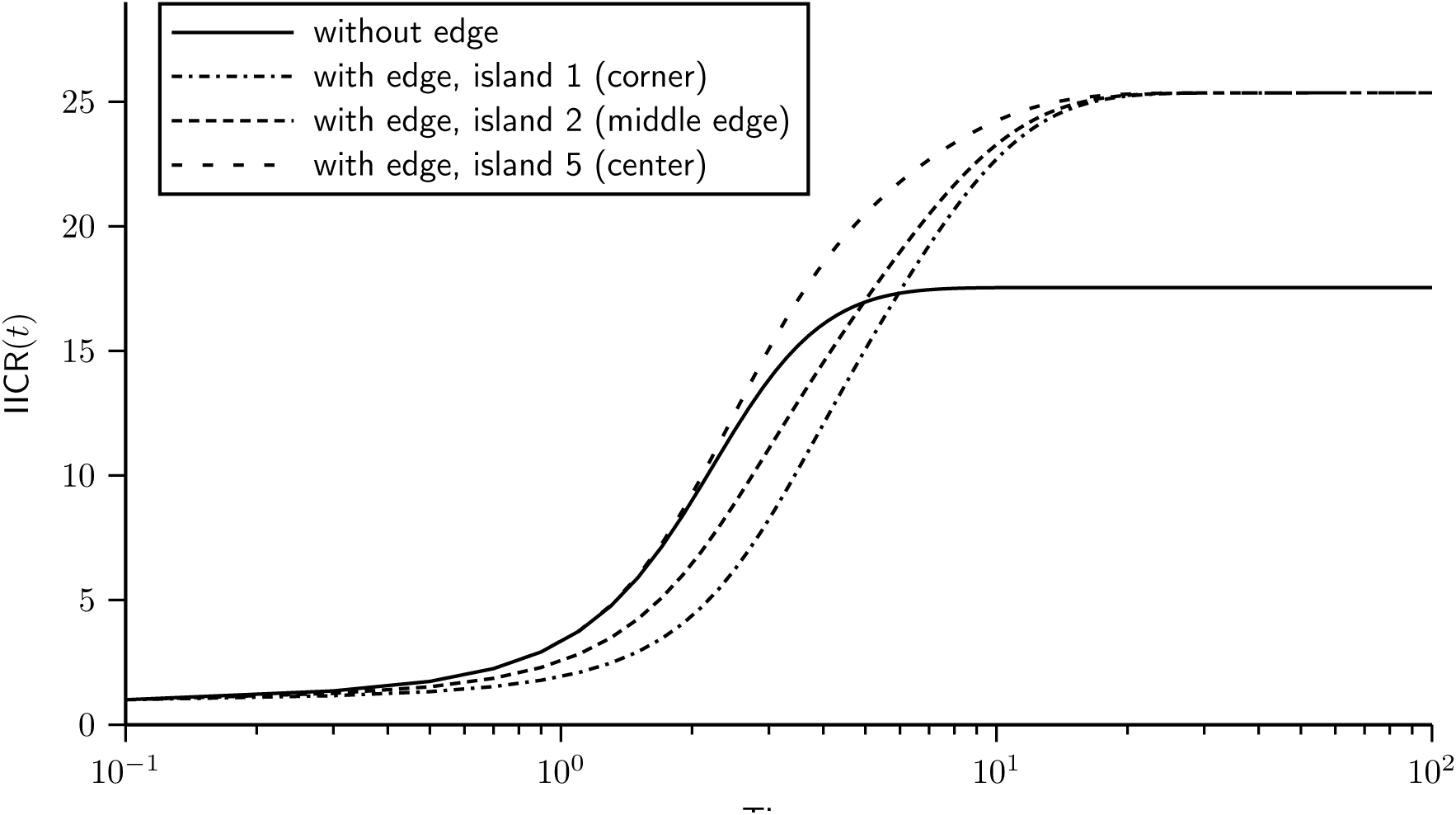
IICR plots for the 2D stepping stone model. Here we assumed a model with 3 × 3 = 9 islands and *M* = 1, with and without edge effect. In the model with edge effect, we plot the three ways to sample two lineages in the same island: in island 1, 3, 7 or 9 (corner), in island 2, 4, 6 or 8 (middle of the edge), and in island 5 (center of the lattice).

### 4.2 Continent-island model

#### 4.2.1 General case

Here we assume a model where the population is divided into *n* demes (one big deme called *continent* and *n* − 1 equally sized demes, smaller than the continent, called *islands*). The continent is connected with the remaining *n* − 1 islands, but the islands are not connected between each other (Figure 2). Therefore, migration can only occur between the continent and the islands, but not between different islands. We order the *n* demes in such a way that the continent is deme number 1, whose (scaled) size is *c*_1_. We denote *c*_2_ the size of the other islands, and *M*_1_/2 the (scaled) migration rate from the continent to each island, and *M*_2_/2 the migration rate from each island to the continent. Condition (1) implies that we have the following constraint:

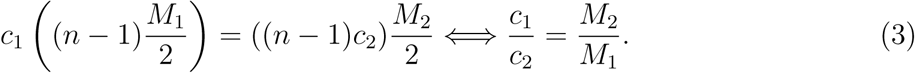

For the case *n* ≥ 3, the symmetry of the model allows us to consider, for a sample of two lineages, only five possible different configurations:

1. Both lineages are in the continent. A coalescence can occur with rate 1/*c*_1_, leading to configuration 5, or any of the two lineages may migrate to one of the *n* − 1 islands, each with rate *M*_1_/2, leading to the second configuration.
2. One lineage is in the continent and the other in an island. There can be no coalescence event, but three different migration events can occur: if the lineage in the island migrates, which arrives at rate *M*_2_/2, this leads to the first configuration. The lineage in the continent can migrate at rate *M*_1_/2, and it can either reach the island where the other lineage is (leading to configuration 4 below) or migrate to a different island (leading to configuration 3 below).
3. The two lineages are in different islands. No coalescence can occur and any of the two lineages can migrate to the continent, each with rate *M*_2_/2, leading to configuration 2.
4. The two lineages are in the same island. Either a coalescence occurs with rate 1/*c*_2_, leading to configuration 5, or a migration event of one of the two lineages to the continent, each with rate *M*_2_/2, leading to configuration 2.
5. The two lineages have coalesced. This is an absorbing state.

If we replace *M*_2_ by *M* and *M*_1_ by *c*_2_*M/c*_1_ in equation (3) and normalise population sizes by fixing *c*_1_ = _1_, then denoting *c*_2_*/c*_1_ = *c*_2_ = *c* we obtain the following transition rate matrix (see Supplementary Materials for details):

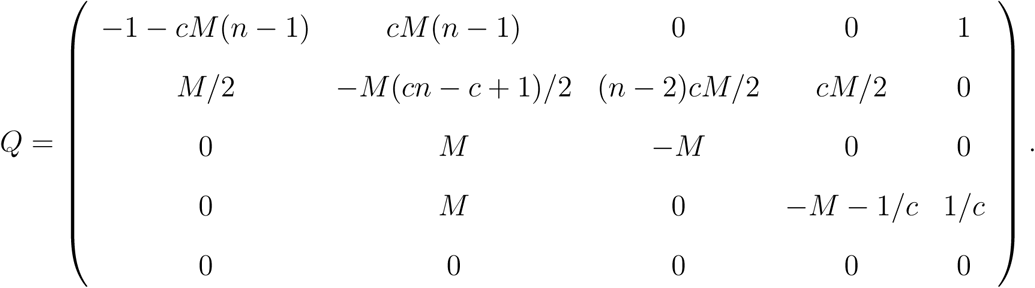

Note that *c* is the ratio between the sizes of the islands and the continent, and that the diagonal entries are obtained by the constraint that the sum over each row is zero.

Figure 4 shows the IICR_s_ and IICR_d_ plots for the different sample configurations for a pairs of genomes in a continent-island model with *n* = 4 (one continent and three islands). As expected from previous work on the IICR (Mazet et al., 2016; Chikhi et al., 2018), first generation hybrid individuals, whose genome is sampled in different demes, exhibit IICR plots which would be interpreted as expansions from an ancient stationary population, even though the total population size is constant. One of the most striking result is that a diploid individual sampled in one of the islands exhibits an IICR that suggests (forward in time) an ancient stationary population which first expanded before being subjected to a significant population decrease. Thus, different individuals will exhibit very different history, not because their populations were subjected to different demographic histories, but because the IICR does not represent the history of a population. It represents the coalescent history of a particular sample in a particular model.

**Figure 4:**
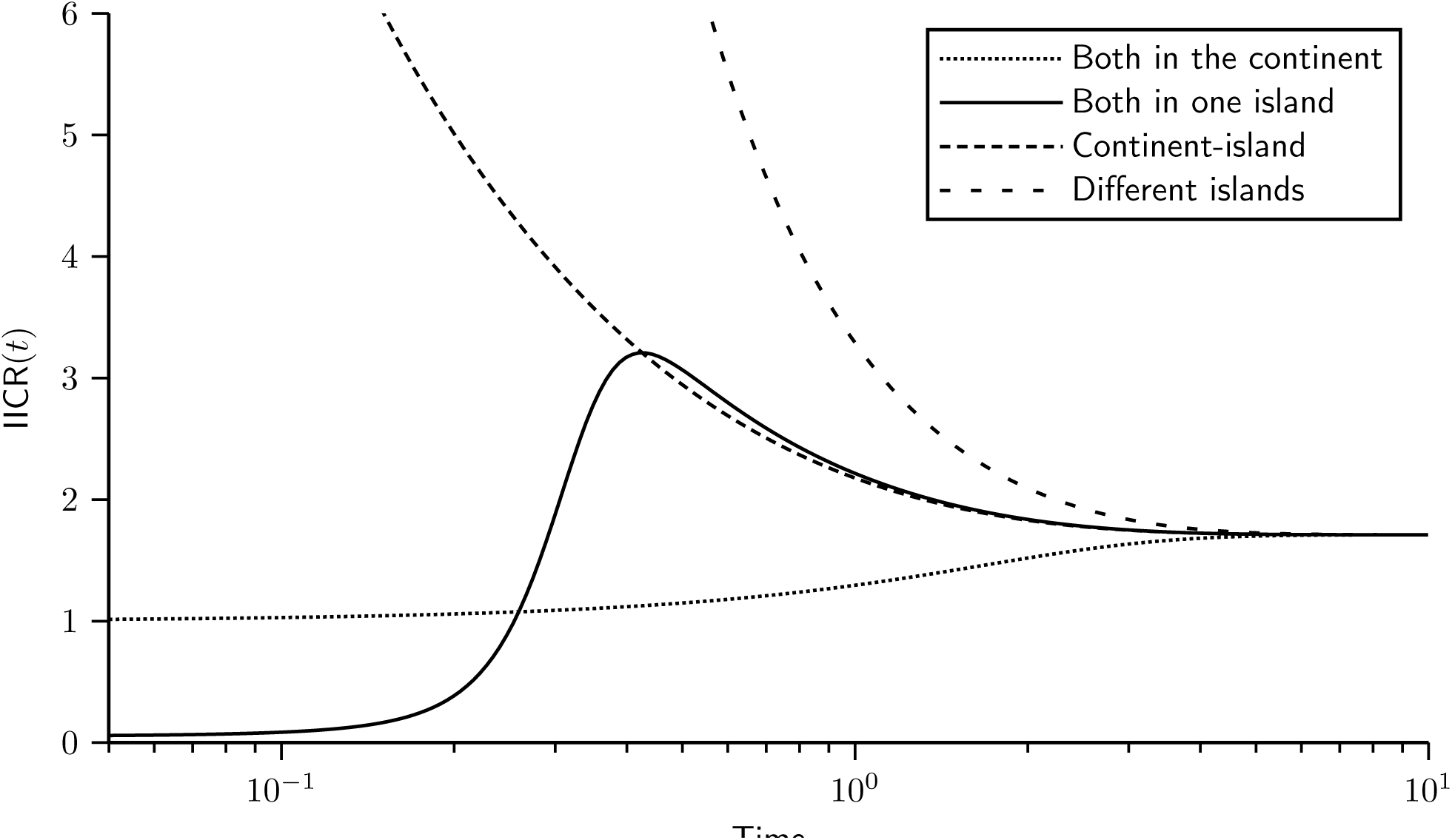
IICR for a continent-island model. We constructed the transition rate matrix for a model with *n* = 4, namely one continent and three same-sized islands. The sizes of the continent and of the islands were set to *c*_1_ = 1 and *c*_2_ = 0.05, respectively. In other words, the continent was 20 times larger than the islands. We set the migration rates to *M*1*/*2 = 0.05, *M*_2_*/*2 = 1 (note that once *M*_1_ is set, *M*_2_ is constrained to keep inward and outward migrant gene numbers equal, as required by equation 1). In this model there are only four types of IICR curves, two IICR_s_ and two IICR_d_. The first two correspond to the cases where we sample the two lineages either in the continent or in one of the islands. The IICR_d_ curves correspond to cases where one gene comes from the continent and the other from an island or when the two genes come from two different islands.

## 5 The Non-Stationary Structured Coalescent (NSSC): constructing the IICR for models with changes in population structure

In this section we extend our work to non-stationary structured (NSS) models under the coalescent and show how the semigroup property can be used to characterise a large family of complex NSS models. The semigroup property allows to compute the probability that a Markov jump process is in a given state at time *t* + Δ*t* by taking into account all its possible states at time *t.* Applied to the structured coalescent, this makes it possible to trace ancestral lineages backward to the MRCA in models where some parameters (*n*, *c_i_, M_ij_*) may change at some time point in the past. In particular, this gives a way to compute (at least numerically) the distribution of coalescence times for a wide family of non-stationary structured models, hence allowing us to introduce and study the NSSC.

### 5.1 Applying the semigroup property to the structured coalescent

Previous sections showed that to any given stationary structured population model corresponds a transition rate matrix, *Q* that can be constructed and used to predict the IICR for a given sample configuration. Assuming that we sample *k* genes in configuration *α,* we call 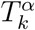 the time to the first coalescence event among these *k* lineages. We also described how the theory of Markov chains allows to compute the probability distribution of 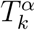 from *Q* using the formula:

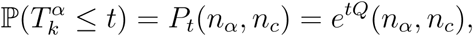

where *n_α_* denotes the index of the configuration *α* and *n_c_* is the number of possible configurations and corresponds to the index of the coalescence configuration.

The matrix *P_t_* (which is the *transition semigroup*) has size *n_c_* × *n_c_* and is obtained by computing the exponential of the matrix *tQ.* The elements of this *n_c_* × *n_c_* matrix are functions of the parameters of the model (*n*, *c_i_, M_ij_*), which are assumed to be constant under the structured coalescent (stationary model). Now, the semigroup property states that for any positive values *t* and *u* we have:

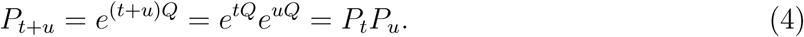

By using the semigroup property, the structured coalescent can be extended to non-stationary models (e.g., models with changes in the size of one or more demes or in the values of gene flow at some point in the past).

For simplicity, we assume here that the number of demes *n* is fixed for a given species. The reason for doing this is that, once we fix the number of genes sampled at the present (*k*) and the number of demes (*n*), the number of possible states or configurations of the Markov process (|*E_k,n_*|) is also fixed and so is the size of the corresponding transition rate matrix. It will be thus straightforward to compute products of matrices, using Equation (4). Keeping *n* constant guarantees that other parameter changes (i.e., *c_i_, M_ij_*) will not modify the state space of the Markov jump process, even if the transition probabilities between these states will change. So, the size of the matrix *P_t_* will always be the same.

Assume that at time *t* = *T* in the past, some of the parameters *M_ij_* or *c_i_* change. This change has no influence on *E_k,n_* and does not affect the evolution of the process between *t* = 0 and *t* = *T.* Denote by *Q*_0_ the transition rate matrix of the Markov chain for 0 ≤ *t* ≤ *T* and *Q*_1_ the corresponding transition rate matrix for *t* ≥ *T.* If we call 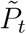 the *transition semigroup* of the Markov chain that models this structured scenario with a demographic change event at time *T,* we can compute 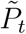 by using the semigroup property as follows:

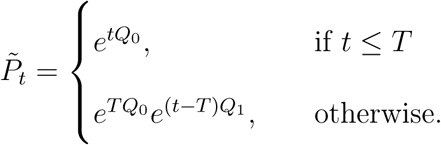

In particular, the distribution of 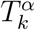, the first coalescence time of *k* genes sampled in configuration *α* under this structured model with a past demographic change event, can be computed by:

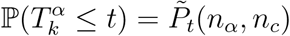

The *pdf* of 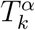 can then be computed by 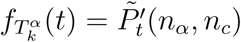, where

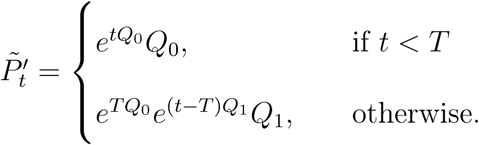

This procedure can be extended to any number of parameter changes, by defining the respective transition rate matrices for each of the time intervals between successive changes in the parameters of the structured model. Thus, the distribution of coalescence times (and the IICR) for structured models in which migration rates and demes sizes can arbitrarily change, can be obtained from the computation of matrix exponentials and matrix products.

Moreover, the NSSC framework allows to compute the IICR for models considering a population split. For example, a model considering one ancestral population that separated into two subpopulation at time *T* can be easily approximated under the NSSC framework. To do this, just set a value of gene flow from the present to time *T.* Then set a gene flow equal to infinity (in practice we use a gene flow high enough so that the two populations behave as a panmictic one) from time *T* to the past. The following section considers a more general model of population split that gives a new perspective to the history of evolution of humans and Neanderthals.

### 5.2 Application: Humans and Neanderthals IICR

In this section we show how a single model (Figure 5) incorporating both humans and Neanderthals as structured species derived from an unknown ancestral *Homo* species that was itself structured, can be used to predict the PSMC plots inferred for humans and Neanderthals (see details below). The IICR for humans and for Neanderthals were predicted using the NSSC framework, assuming that one diploid was sampled in a human deme and another in a Neanderthal deme. Following the approach used by Chikhi et al. (2018) we also computed the IICR using *T*_2_ values simulated with Hudson’s *ms* software for the same demographic scenario. Finally, the PSMC plots inferred from real data are also plotted in the same panel for comparison. As an additional validation step we also plot in panel b the PSMC inferred from genomic data simulated with *ms* (i.e., DNA sequences rather the *T*_2_ values) for the same scenario together with the PSMC from the real sequences.

**Figure 5:**
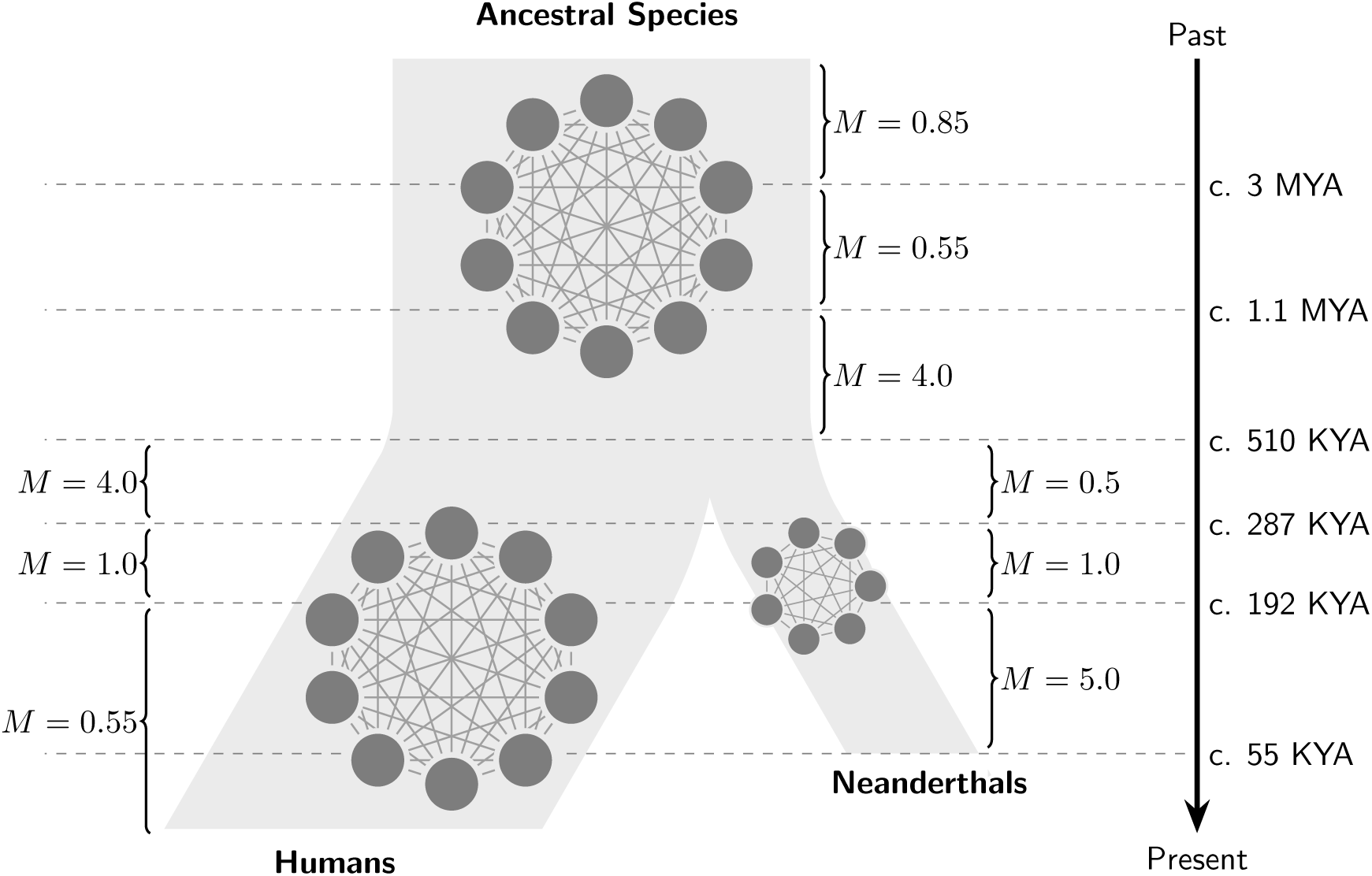
Hypothetical scenario presenting humans and Neanderthals as structured species derived from and unknown *Homo* species that was itself structured. The times at which gene flow (*M*) changed are indicated by horizontal lines.

In the proposed scenario (see Figure 5), humans and Neanderthals descend from a *Homo* species that was structured in ten interconnected demes, as in Mazet et al. (2016), and whose connectivity changed around 3 million years ago (MYA) when the migration rate *M* = 4*Nm* decreased from 0.85 to 0.55. Then, around 1.1 MYA, *M* increased significantly from 0.55 to 4. The following period of reasonably high connectivity (*M* = 4 corresponds to an *F_st_* of 0.11 across the whole species) was maintained in the lineage that led to humans until 0.287 MYA whereas a significant change occurred when Neanderthals split from that common lineage, some time about 0.51 MYA. Our model suggests that to fit the estimated Neanderthal PSMC results the original Neanderthals are the result of a “sub-sampling” or split from human demes (*n* = 7 demes in our model). These new Neanderthal demes were around 16% of the size of human demes. At the same time (0.51 MYA) *M* decreased from 4 to 0.5 in the Neanderthal lineage whereas, as noted above, it remained constant in humans. In the case of Neanderthals, the reduction is surprisingly close to the level of connectivity of the ancestral species (between 3 and 1.1 MYA). It is as if archaic Neanderthals were a group of small demes that derived from human demes and that had gone back to an ancestral low connectivity state. Neanderthals stayed in that low connectivity state until 287 KYA. One striking result is that a simultaneous change is observed at that time in humans and Neanderthals, and that it is now in the opposite direction. Whereas gene flow started to decrease in humans, from *M* = 4 to *M* = 1, it doubles in Neanderthals from *M* = 0.5 to *M* = 1. Then, around 192 KYA, gene flow increases to *M* = 5 in Neanderthals and decreases to *M* = 0.55 in humans. It is as if in a period of 100 KY Neanderthals’ gene flow had increased 10-fold, perhaps as a consequence of a geographic contraction. Humans on the other hand appear to have maintained a low connectivity until the Neolithic as discussed in Mazet et al. (2016). Assuming a mutation rate per generation equal to 1.25 × 10^−8^, the proposed scenario is consistent with a deme size of 1276 for humans and a deme size of 200 for Neanderthals. Note that under this scenario, deme sizes remain constant and the PSMC patterns can be explained only by changes in connectivity. Note also that in this figure, we did not simulate the Neolithic expansion, which is why the human IICR and PSMC plots continue to decrease to the local deme size in the recent past, as explained in Mazet et al. (2016) and Chikhi et al. (2018).

If we trace the theoretical IICR corresponding to the scenario described above, we can see that it is similar to the PSMC plots obtained from real human and Neanderthal data (Figure 6). Moreover, we simulated 40 full genome length (i.e., 3 GB) sequences with *ms* under the proposed scenario. The first 20 corresponded to a genome sampled in a human deme and the last 20 corresponded to a genome sampled in a Neanderthal deme. We then applied the PSMC to each of these simulated sequences and compared the results with the PSMC plots obtained from real data (Figure 6).

**Figure 6:**
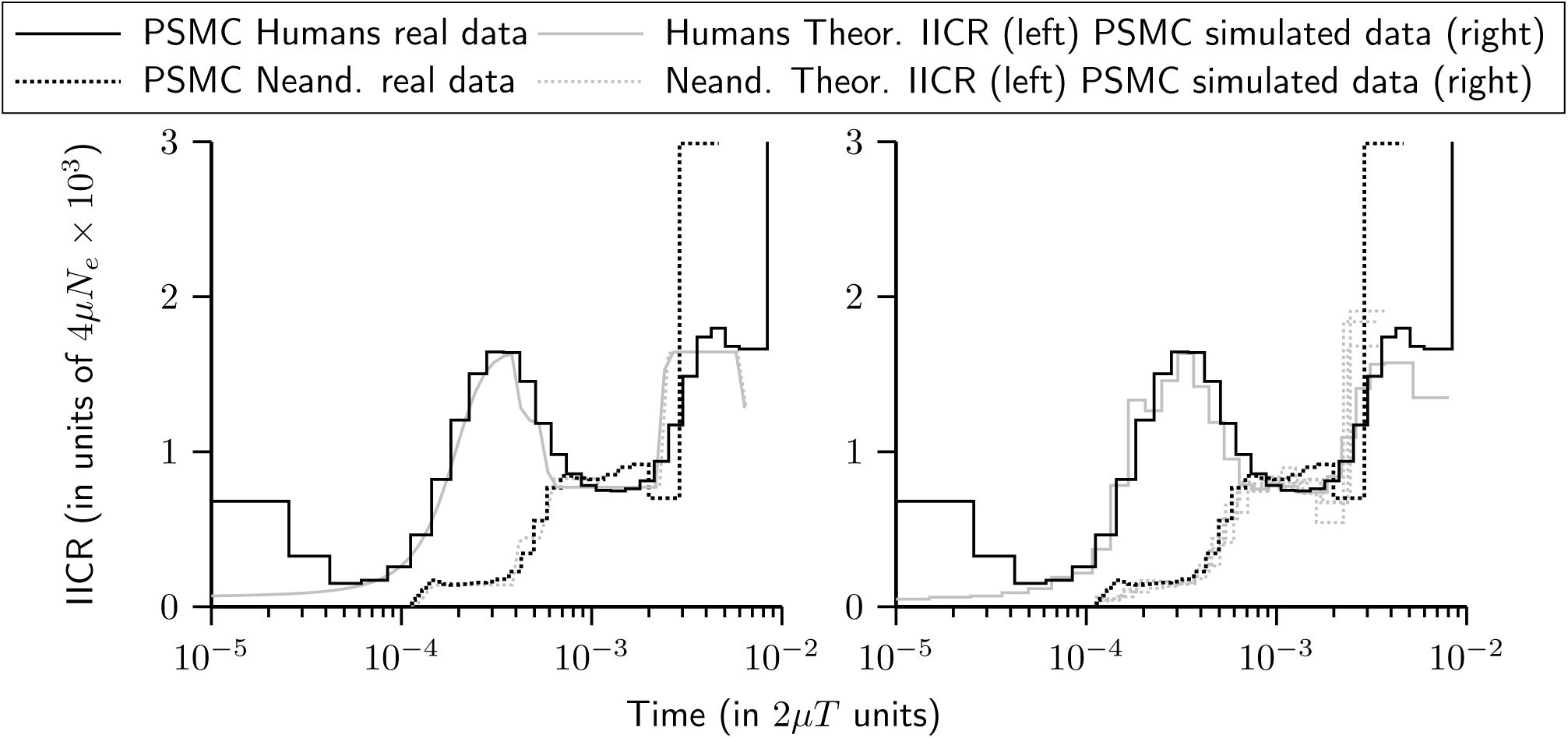
IICR and PSMC plots for humans and Neanderthals. The PSMC plots obtained from real human and Neanderthal sequences are similar to the theoretical IICR (left panel) corresponding to the proposed scenario. Also, they are similar to the PSMC plots obtained from sequence data simulated under the proposed scenario (right panel).

It is worth stressing that the absolute dates presented here should be taken with a grain of salt since they depend on various parameters which we took from previous studies. In Mazet et al. (2016) and Chikhi et al. (2018) we used the mutation rates of Li and Durbin (2011) but here we used the values of Prufer et al. (2014) to be able to compare our IICR results to the PSMC results obtained by the latter study. This explains why several dates are shifted compared to those of Mazet et al. (2016).

Altogether, these results show that the scenario proposed explains the skyline plots obtained by PSMC from real data. It is thus possible to construct a scenario in which humans and Neanderthals are structured and descend from a common ancestral species that was also structured. PSMC plots are usually interpreted in terms of population size change. However, this scenario explains PSMC plots without any change in population size in humans, and with a split, disconnection and deme size reduction in Neanderthals. The scenario, however, requires neither gene flow nor admixture between humans and Neanderthals. The simple fact of sampling diploids in different demes (humans or Neanderthals) generates the very different PSMC plots inferred for humans and Neanderthals.

## 6 Discussion and perspectives

### 6.1 The NSSC as an extension of the structured coalescent

The theoretical framework presented in this study is closely related to Herbots’ works (Herbots, 1994; Wilkinson-Herbots, 1998), who introduced the use of transition rate matrices for studying structured models and computed the coefficients of the transition rate matrix for many stationary models. Here we extended the existing theory to non-stationary structured models. This can impact future population genetic studies in several important ways. The NSSC framework gives a theoretical way for computing the *cdf* and the *pdf* of *T*_2_ under a wide family of models of structured population. It also includes a natural way of incorporating past demographic events (i.e., changes in deme sizes and/or in gene flow) into models of population structured. Currently, most of the population genetic studies either assume panmixia and try to infer past changes in population size or consider that population is structured and infer parameters related to the structure without taking into account the changes in the population size. The NSSC framework developed here is original because it allows to combine changes in population structure and size into the same model. Allowing to incorporate past demographic events into a model considering population structure is a step forward that may help to disentangle the confounding effects of structure on methods used to reconstruct demographic history that has been pointed by previous studies (Chikhi et al., 2010; Heller et al., 2013).

Moreover, given that theoretical distribution of *T*_2_ is known, we can use numerical approximations to compute corresponding IICR curves with much lower computational time than the simulation based approach used in Chikhi et al. (2018). This gives a very quick way of testing alternative scenarios and also lays the theoretical bases to implement an inferential framework using the IICR computed from genomic data by methods like PSMC Li and Durbin (2011) or MSMC (Schiffels and Durbin, 2013). However, the construction of a such inferential process as well as the corresponding validations for simple and complex models would need a full and independent study.

We would like to stress that the theoretical arguments that guarantee the convergence of the discrete-time process described in 2.1 to a continuous-time Markov process lay on the assumption given in equation 1 (see Herbots (1994) for details). However, some authors have proposed methods based on the same approximation to a continuous-time Markov process without taking condition 1 into account (Notohara, 1990; Costa and Wilkinson-Herbots, 2017). Moreover, simulation software like the popular *ms* (Hudson, 2002) do not necessarily make this hypothesis when dealing with structured models. The question of whether the hypothesis given in 1 is crucial or can be removed without affecting the convergence to the continuous-time Markov process is beyond the objectives of this work and deserves an independent study too.

### 6.2 Humans, Neanderthals, and genomic story-telling

While the scenario proposed in 5.2 should not be taken at face value, some hypothesis and interpretations based on a such scenario may be interesting. This scenario suggests that one major event dated around 290 KYA induced a change in connectivity that was simultaneous in humans and Neanderthals. In this sense, we identify a striking consistency across the two species. One interpretation could be that the two *Homo* species responded to the same environmental change, around 290 KYA, one species (Neanderthals) with an increase in connectivity as a possible consequence of a spatial contraction and the other (humans) with a decrease in gene flow between populations, as a possible consequence of geographic expansion towards new territories. Another interpretation would be that only one of the species (most likely humans), reacted to a major environmental change or experienced a major behavioural change, that are both yet to be identified. This change in distribution may have led to a change in the interactions humans had with Neanderthals perhaps as a consequence of a human geographical expansion. This could have led the Neanderthals to contract. By doing so, Neanderthal populations that used to be little connected started to interact more and behave increasingly like a panmictic population, hence reducing the apparent *N_e_* (or more precisely reducing the IICR). For reasons that we can only speculate on, Neanderthals went extinct not because they became separated and isolated, but rather as a consequence of a likely reduction of their distribution which led to an increase of gene flow after long periods during which they survived as small isolated populations.

One should be very careful at this stage as there is not much Neanderthals’ genomic data available that could make possible to infer the PSMC for other individuals and determine if there is a signature of spatial structure. Here, we focused on *n*-island (i.e., non-spatial) models, even though we have noted in Chikhi et al. (2018) that spatial models will likely be necessary to explain the diversity of human PSMC plots. We stress however that the proposed structured model provides a new and fundamentally different outlook on Neanderthals extinction. Our model explains the decrease in the Neanderthal PSMC plots, not as a decrease in population size but rather as a result of decreased isolation of Neanderthal populations, and as a consequence of the properties of the IICR in structured models. Indeed, the “humps and bumps“of IICR plots (Chikhi et al., 2018) can be caused by changes in connectivity or by a constitutive property of the IICR (Mazet et al., 2016; Chikhi et al., 2018).

While the presented scenario does not aim to explain all the complexity of human and Neanderthal evolution it explains genomic patterns that are currently not explained by several existing admixture models. For instance, Chikhi et al. (2018) used coalescent simulations of *T*_2_ values to compute the IICR for several models of population structure, and applied their simulation-based approach to the admixture and ancient structure models of Yang et al. (2012). They found that none of the models used by Yang et al. (2012) could explain the PSMC plots of humans and Neanderthals even though some admixture models could explain a modified allele frequency spectrum better than models without admixture. Here we proposed a new scenario that can explain the PSMC plots of Neanderthals and humans and is thus consistent with a no admixture history between humans and Neanderthals. This model is in agreement with Eriksson and Manica (2012) who argued that the D-statistic used to quantify Neanderthal admixture is influenced by population structure.

Similarly, Kuhlwilm et al. (2016) used a model with splitting populations to represent the evolution of humans, Neanderthals and Denisovans. Their model was not inferred from the data but rather chosen *a priori* and probably on the basis of beliefs (or knowledge) that the authors had gathered. While they did carry out several validation steps, the model was not inferred from the data. Based on our understanding of the IICR in structured models Mazet et al. (2016); Chikhi et al. (2018), it seems very unlikely that their model could explain the known PSMC curves of humans and Neanderthals. For instance their model assumes constant population sizes and ignores gene flow one of which at least is typically necessary to generate humps and bumps in IICR plots Mazet et al. (2016); Chikhi et al. (2018).

The fact that we mainly used models without changes in population size does not mean that we believe that there were no changes in deme size in the history of most species including humans or Neanderthals. It however means that such changes are not always necessary to explain the data and that changes in connectivity should be better integrated in our understanding of the recent evolution of species Chikhi et al. (2010); Mazet et al. (2016); Chikhi et al. (2018). Mazet et al. (2016); Chikhi et al. (2018) showed how different individuals from the same species can exhibit very different “demographic histories” simply because they or their genes were sampled in different locations of a structured population.

Changes in connectivity in a complex splitting model produce complex genomic patterns that cannot be easily interpreted. By using the IICR and the NSSC we were able to re-interpret human and Neanderthal evolution, while stressing that it is only one of probably many possible interpretations.

The structured scenario used here for humans and Neanderthals ignores spatial structure but Chikhi et al. (2018) noted that to understand human evolution, spatial models such as stepping stone models would probably be necessary to explain the variability observed in human PSMC plots. For Neanderthals similar claims cannot be made yet since only one Neanderthal PSMC plot has been published to date. In our model, when Neanderthals split from the common ancestral species, they have much smaller demes than humans and these demes are less connected. It is interesting to note that a recent genomic study by Rogers et al. (2017) suggested that Neanderthals were probably distributed in small and isolated demes. Our results are thus consistent with that idea. We note though that there are significant differences. In our model, Neanderthals saw a significant increase in gene flow around 290 KYA (maybe more recently depending on the mutation rate) and again around 190 KYA.

The fact that two sets of independent models can explain humans and Neanderthal PSMC plots suggests that admixture between humans and Neanderthals is not necessary to explain human or Neanderthals PSMC plots. We thus conclude with Chikhi et al. (2018) that claims of admixture may be weaker than usually believed, even if we must also conclude that admixture cannot be excluded today.

Beyond humans and Neanderthals, the NSSC modelling presented here should now be developed as a full inferential tool to identify quickly and efficiently models that can, and models that cannot, explain known genomic features. The transition rate matrices approach can make the computation of the IICR extremely efficient. This suggests that the IICR can be computed for various models and compared to observed PSMC plots. It can thus be used as a summary of genomic data and estimated with the PSMC and MSMC methods, as suggested by Chikhi et al. (2018) to exclude models or identify the best models.

### 6.3 Increasing the sample size to more than two sequences

The Markov process approach used in sections 2 and 5 allows to trace back ancestral lineages coming from a sample of arbitrary size. This means that we can compute the distribution of the first coalescence event in a sample of *k* genes (denoted *T_k_*) for *k* ≥ 2. Thus, it is theoretically possible under the NSSC framework to obtain statistical properties of the underlying genealogical tree for samples of size *k.* However, in this study we mainly focused on the IICR as defined by Mazet et al. (2016) for *T*_2_. The reason for this is that when *k* ≥ 3 the number of states to consider in the Markov process becomes very large and so does the corresponding transition rate matrix. It becomes messy to enumerate all the states and to construct the corresponding transition rate matrix. Moreover, the computation of the matrix exponential becomes intractable under the classic numerical methods (Moler and Loan, 2003). Some optimisations need to be done taking advantage of the particular structure of the matrices associated to the NSSC framework. Also there is a need for a clear algorithm enumerating all the possible states when tracing back more than two ancestral lineages to the MRCA. It may also be possible to construct a “reduced” transition rate matrix instead of the one if there are “symmetries” in the model. For instance, the *n*-island model is highly symmetrical (all islands have the same size and migration rates are identical between all islands). The advantage of using symmetries is that it significantly reduces the size of the transition rate matrix and computation time but this idea will not be viable for all structured models.

In conclusion, one of the great challenges of population genetics inference is to identify the structured models that could explain existing genomic data. Until now the choices of structured models has been to a large extent arbitrary. The NSSC modelling framework proposed here may be a powerful and promising way to overcome that challenge, and perhaps reduce arbitrariness and some level of story-telling that has often plagued human evolution discourse. All scripts used to carry out the simulations or analyse the data will be made available upon publication of the manuscript.

## 7 Acknowledgements

We would like to thank Didier Pinchon and Armando Arredondo for valuable comments and contributions to the manuscript. We would also like to thank Josue Corujo for productive discussions about Markov chains theory and its applications to the structured coalescent. This research was funded through the 2015-2016 BiodivERsA COFUND call for research proposals, with the national funders ANR (ANR-16-EBI3-0014), FCT (Biodiversa/0003/2015) and PT-DLR (01LC1617A). This work was also funded by the PHC PESSOA 2016/2017 program (ref. 354652NK) between Portugal and France. This work was supported by the French Laboratory of Excellence project “TULIP” (ANR-10-LABX-41; ANR-11-IDEX-0002-02) and the the LIA BEEG-B (Laboratoire International Associé — Bioinformatics, Ecology, Evolution, Genomics and Behaviour) between the CNRS and IGC. We also acknowledge an Investissement d’Avenir grant of the Agence Nationale de la Recherche (CEBA : ANR-10-LABX-25-01). We are grateful to the genotoul bioinformatics platform Toulouse Midi-Pyrenees (Bioinfo Genotoul) for providing help and/or computing and/or storage resources.

## Supplementary Information

### 1 General algorithm for the construction of the transition rate matrix for two lineages

We give a general algorithm that can be used to construct the transition rate matrix of a given model. The first step is to explicitly order all the demes. Then, given the number *n* of (ordered) demes the set of all possible configuration for *k* = 2 lineages is:

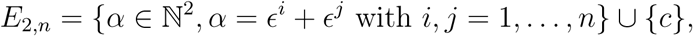

We take the inverse lexicographical order on *E*_2*,n*_. Define *ϕ* as a function from *E*_2*,n*_ to {1, 2, …, |*E*_2,*n*_|} such that *ϕ*(*α*) is the index of *α* according to the inverse lexicographical order. Then *ϕ*^−1^ is the inverse of *ϕ* and *ϕ*^−1^(*i*) gives the element of *E*_2,*n*_ which is at position *i* according the inverse lexicographical order.

Once the function *ϕ* is defined and we have the values of *C* = (*c*_1_,…*, c_n_*) (the size of the demes) and *M_ij_* (the migration matrix), we can use the following algorithm to construct the transition rate matrix *Q:*

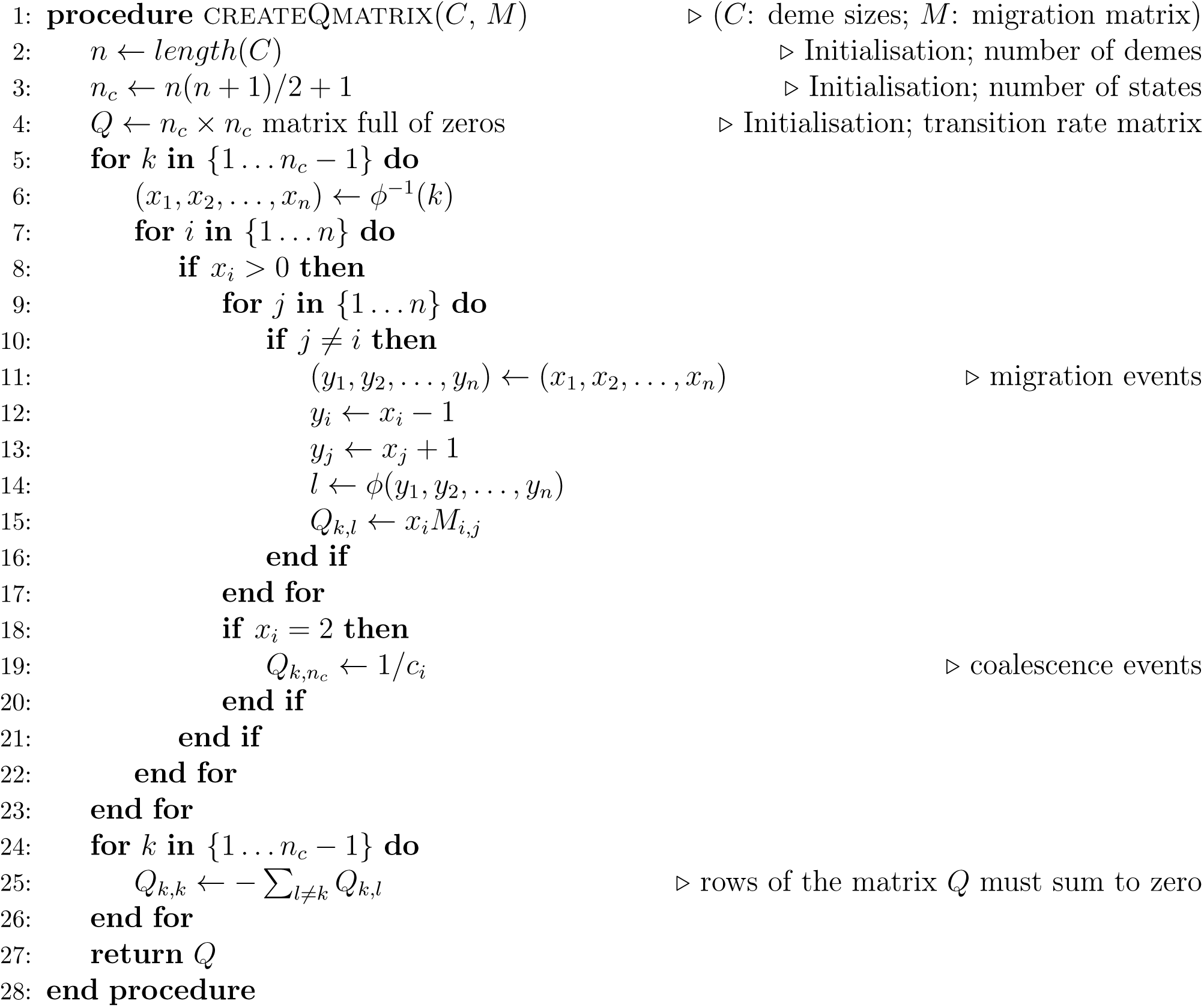

### 2 Constructing the IICR for stationary models. Examples: stepping stone and continent-island

We now apply the framework and algorithm described above to some stationary models. By a stationary model we understand a structured model in which the parameters (i.e., number of demes, sizes of demes and gene flow) remain constant over time. To our knowledge, there is no analytical expression for the distribution of the coalescence time *T*_2_ under these models. For some of them it is possible to find a simplified transition rate matrix using some symmetries. In those case we give the corresponding transition rate matrix *Q* that can be used to compute numerically the distribution of *T*_2_ and the IICR. In other cases it is not possible to get a simplified version of *Q* and we used the algorithm given in section 1 to obtain the IICR.

#### 2.1 stepping-stone models

Stepping stone models (Kimura, 1953; Malécot, 1948) assume that the demes are located at the nodes of a regular lattice in one or two dimensions (hereafter 1D and 2D stepping stone models). Each deme can have up to four neighbours and migration events are only possible between neighbouring demes. These models incorporate space, and are thus thought to be more realistic than the *n*-island model described above, which implicitly assumes that migration is as likely between neighbours as it is between distant islands. The border demes can either be connected with each other, hence forming a torus, or can behave as bouncing borders (Figure S1). In some models the bouncing borders migrants are assumed to stay in their deme, whereas in other models they are distributed among the demes to which their deme is connected.

We will distinguish two cases:

1. Without edges: One dimension (1D circular stepping stone) and two dimensions (2D torus stepping stone). They are more symmetric since all the migration rates are equal.
2. With edges: 1D and 2D stepping stone. Islands located on the edges and in the corners have fewer neighbours than islands in the middle of the lattice. In order to maintain simplicity and symmetry, the same migration rate is taken between each pair of islands. This implicitly assumes that migrants trying to migrate “outside” are bouncing back to their deme of origin. As we will see there are still more parameters in the model, and the corresponding transition rate matrices are more complex.

We will give an example of each of the four combinations: one or two dimensions, and with and without edge effects.

##### 2.1.1 Circular 1D stepping-stone model

Here we assume that the population is divided into *n* (*n* ≥ 2) equal-sized islands which are located on a circle (Figure S2). Each island thus receives immigrants coming only from its two neighbours.

With the notations of the main manuscript, ∀*i* = 1… *n* we set *c_i_* = 1, *M_i_* = *M,* and *M_ij_* = *M*/2 if |*i* − *j*| = 1 or |*i* − *j*| = *n* − 1, *M_ij_* = 0 if not.

The symmetry of the model allows us to consider that the configuration of a sample of two lineages depends only on their distance *d,* defined as the number of islands separating them, *d* ranges from 0 to ⌊*n*/2⌋ (⌊*x*⌋ is the largest integer not larger than *x*), that is, ⌊*n*/2⌋ + 1 different values.

The corresponding matrix *Q* is then of size ⌊*n*/2⌋ + 2, the last configuration corresponding to the coalescence event, which can occur only if both lineages are in the same island. When there are five demes (*n* = 5), then we have ⌊*n*/2⌋ = 2, the simplified transition rate matrix *Q* has thus 4 rows and columns:

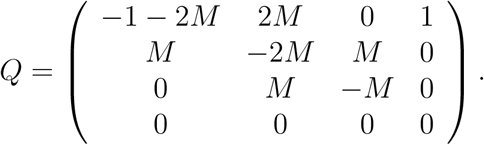

The first row represents the transitions away from the configuration in which both lineages are in the same island. They coalesce with rate 1*/c_i_* = 1. Each lineage can migrate with rate *M/2* towards any of the two neighbouring islands. Any of these migrations will lead to a configuration in which both lineages are in a pair of islands distant of 1 unit (this is the second configuration that we consider).

From this second configuration (corresponding to the second row), no coalescence can occur and each lineage can only migrate to the next island, leading to two possible configurations. Either the migration brings them back on the same island (and we are back to the first configuration with rate *M*/2) or one of them migrates to the next island hence increasing the distance between them by one unit to 2 units (this is the third configuration). Since there are *n* = 5 islands there cannot be a distance greater than two (islands 2 and 5 or islands 1 and 4 are only 2 units distant) and we have thus all possible configurations of the simplified matrix *Q.* Also, since *n* = 5 is odd, migration events from this third configuration can only lead to configurations that are identical to itself or to the second one (with rate *M*/2) (see Figure S2). Some IICR corresponding to the circular stepping stone are shown in Figures S3, S4 and S5.

When there are six demes (*n* = 6), then we have ⌊*n*/2⌋ = 3, the simplified matrix *Q* has thus 5 rows and columns:

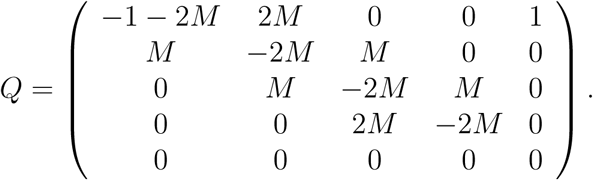

The only difference with the previous example is the fourth configuration, which corresponds to the largest distance of 3 units. From that configuration all migration events necessary lead to the third configuration (corresponding to a distance of 2).

##### 2.1.2 1D stepping-stone model with bouncing edges

Here we consider the edge effects since the two islands located at the extremes of the 1D stepping stone have only one neighbour. The population is divided into *n* (*n* ≥ 2) equal-sized island (see Figure S6).

Keeping the same notations, ∀*_i_* = 1…*n* we set *c_i_* = 1, and 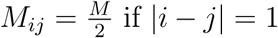, *M_ij_* = 0 if not.

Since there are fewer symmetries than in the circular model, there are significantly more possible configurations in the simplified transition rate matrix *Q* and we now have to take into account the distance between the two lineages, and the distance from the edge of the linear stepping stone.

The general case can be analysed using combinatorics approaches but this will not be presented here and we will simply give the results for *n* = 4. Even in this case the simplified version of the transition rate matrix *Q* has as many as seven rows and seven columns. If we denote by (*i, j*) the configuration when one lineage is in island *i* and the other in island *j,* with *i, j* = 1…4, and given the central symmetry of the model, we can enumerate the configurations as follows :

1. (1, 1) which is the same as (4,4)
2. (1, 2) which is the same as (3,4)
3. (1, 3) which is the same as (2,4)
4. (1,4)
5. (2,2) which is the same as (3,3)
6. (2,3)
7. coalescence *c*

This allows us to construct the corresponding matrix *Q:*

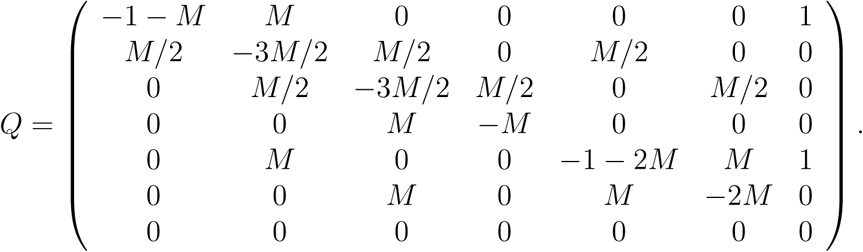

The IICR corresponding to a 1D stepping stone with bouncing edges is shown in Figure S7.

##### 2.1.3 2D stepping stone models with and without edges

For the 2D stepping stone model, we set, ∀*i*, *j* = 1, … *, n, c_i_* = 1 and *M_ij_* = *M/*4 if islands *i* and *j* are neighbours, and *M_ij_* = 0 otherwise. The difference between the models with and without edges used here is thus in the way neighbours are defined. In the model with borders the four corner islands have only two neighbours, the islands on the edges of the lattice have three, and the others have four neighbours (see Figure S1). In the 2D stepping stone model, we computed the corresponding transition rate matrix from the migration matrix of the model using the algorithm given in section 1.

Figure S8 shows the IICR_s_ (two haploid genomes sampled in the same deme, or one diploid genome), for a 3 × 3 stepping stone model with and without borders (Figure S1). In the latter case (no borders), all demes are statistically identical, and there can thus be only one IICR_s_ plot. In the model with borders, there are three possible ways to sample a diploid individual, and three IICR_s_ are plotted. This figure confirms the results of Chikhi et al. (2018) by showing that the IICR_s_ plots for a stepping stone are also S-shaped. They all start in the recent past at a value equal to the deme size and converge in the ancient past towards the same plateau. However, it is remarkable that they differ in the trajectory from the present to the plateau value, depending on the location of the deme (corner, border or centre). These results thus confirm that in a stepping stone model, two diploid individuals sampled in different demes (i.e., geographical regions) will both exhibit signals of population decrease that will be different even though the population size was constant and they both belonged to the same structured model (Chikhi et al., 2018).

#### 2.2 Continent-island model

##### 2.2.1 General case

Here we assume a model where the population is divided into *n* demes (one big deme called *continent* and *n* − 1 equally sized demes, smaller than the continent, called *islands*). The continent is connected with the remaining *n* − 1 islands, but the islands are not connected between each other (Figure S1). Therefore, migration can only occur between the continent and the islands, but not between different islands. We order the *n* demes in such a way that the continent is deme number 1, whose (scaled) size is *c*_1_. We denote *c*_2_ the size of the other islands, and *M*_1_/2 the (scaled) migration rate from the continent to each island, and *M*_2_/2 the migration rate from each island to the continent. Recall that we have the following condition:

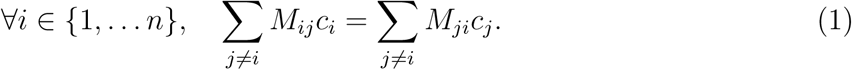

This implies the following constraint:

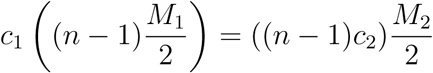

and thus

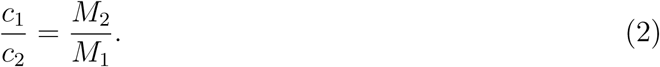

We can thus construct, for the case when *n* ≥ 3, the following 5 × 5 transition rate matrix for a sample of size two (remembering that diagonal terms are obtained such that the sum of the the terms is zero over each row) :

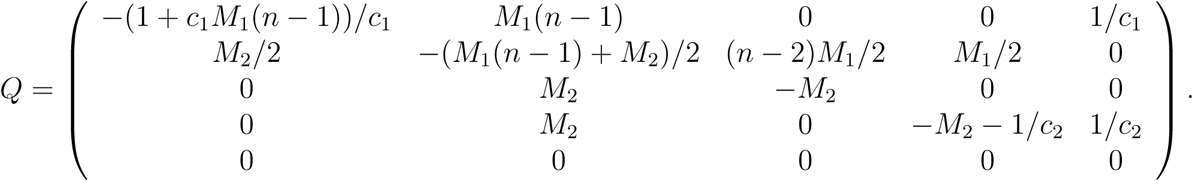

If we replace *M*_2_ by *M* in equation (2) we have *M*_1_ = *c*_2_*M/c*_1_. Then, we normalise population sizes by fixing *c*_1_ = 1. Denoting *c*_2_*/c*_1_ = *c*_2_ by *c,* we obtain the following transition rate matrix:

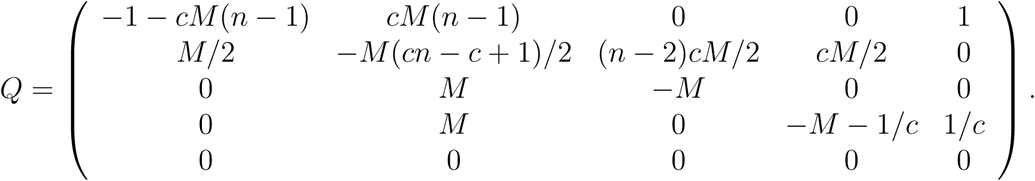

Figure S9 shows the IICR_s_ and IICR_d_ plots for the different sample configurations for a pair of genomes in a continent-island model with *n* = 4 (one continent and three islands). As expected from previous work on the IICR (Mazet et al., 2016; Chikhi et al., 2018), first generation hybrid individuals, whose genome is sampled in different demes, exhibit IICR plots which would be interpreted as expansions from an ancient stationary population, even though the total population size is constant. One of the most striking result is that a diploid individual sampled in one of the islands exhibits an IICR that suggests (forward in time) an ancient stationary population which first expanded before being subjected to a significant population decrease. Thus, different individuals will exhibit very different history, not because their populations were subjected to different demographic histories, but because the IICR does not represent the history of a population. It represents the coalescent history of a particular sample in a particular model.

##### 2.2.2 Particular case: only one continent and one island

If we focus on the particular case where there is only one continent and one island (i.e. *n* = 2), then configuration 3 in the case *n* ≥ 3 does not exist anymore. We thus obtain the following 4 × 4 transition rate matrix:

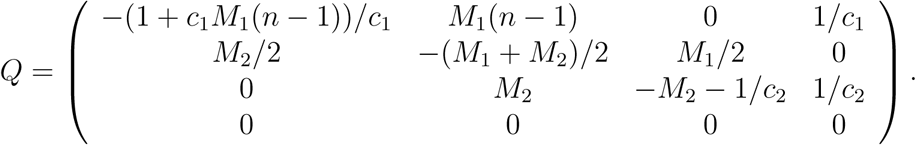

When we replace *M*_2_ by *M* and *c*_1_ by 1 as above, we get:

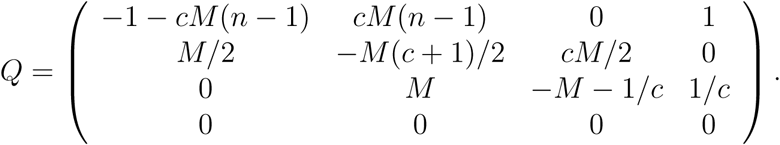

---

**Figure S1:**
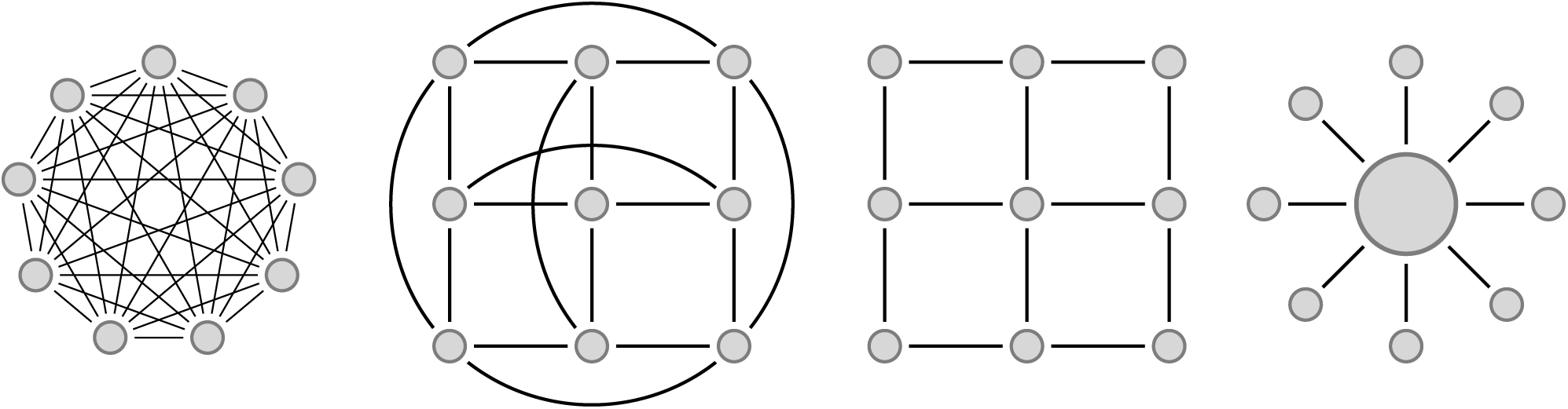
Diagrams for commonly used structured models. From left to right: *n*-islands, torus 2D stepping stone, 2D stepping stone and continent-island model.

**Figure S2:**
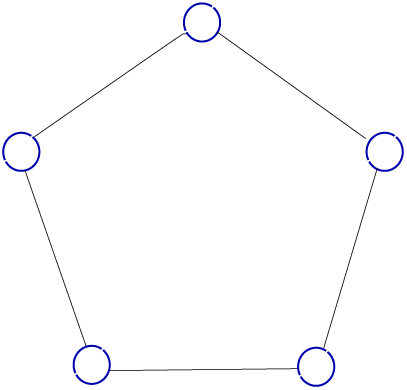
1D circular stepping stone with 5 islands

**Figure S3:**
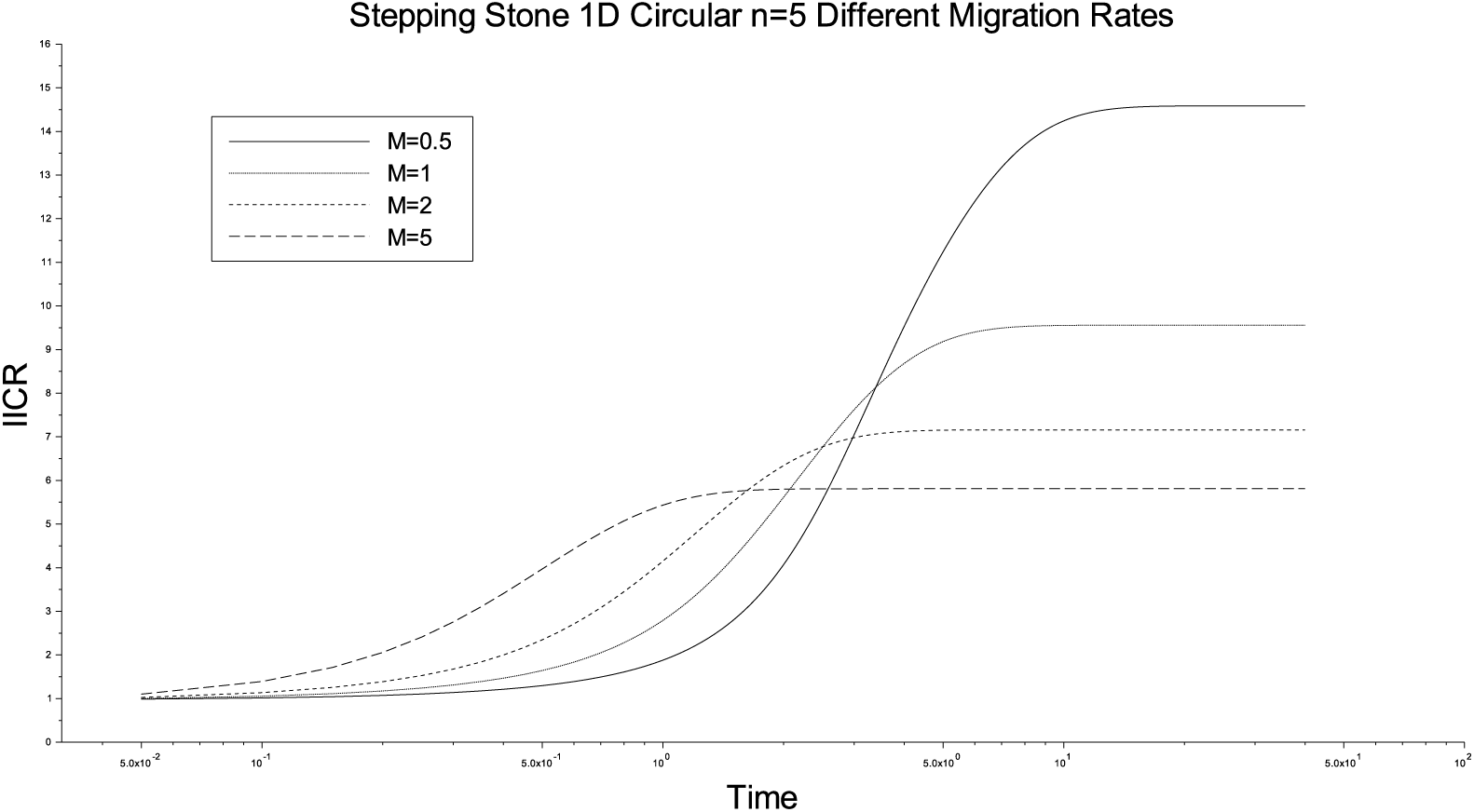
1D circular stepping stone, *n* = 5, different values of *M*

**Figure S4:**
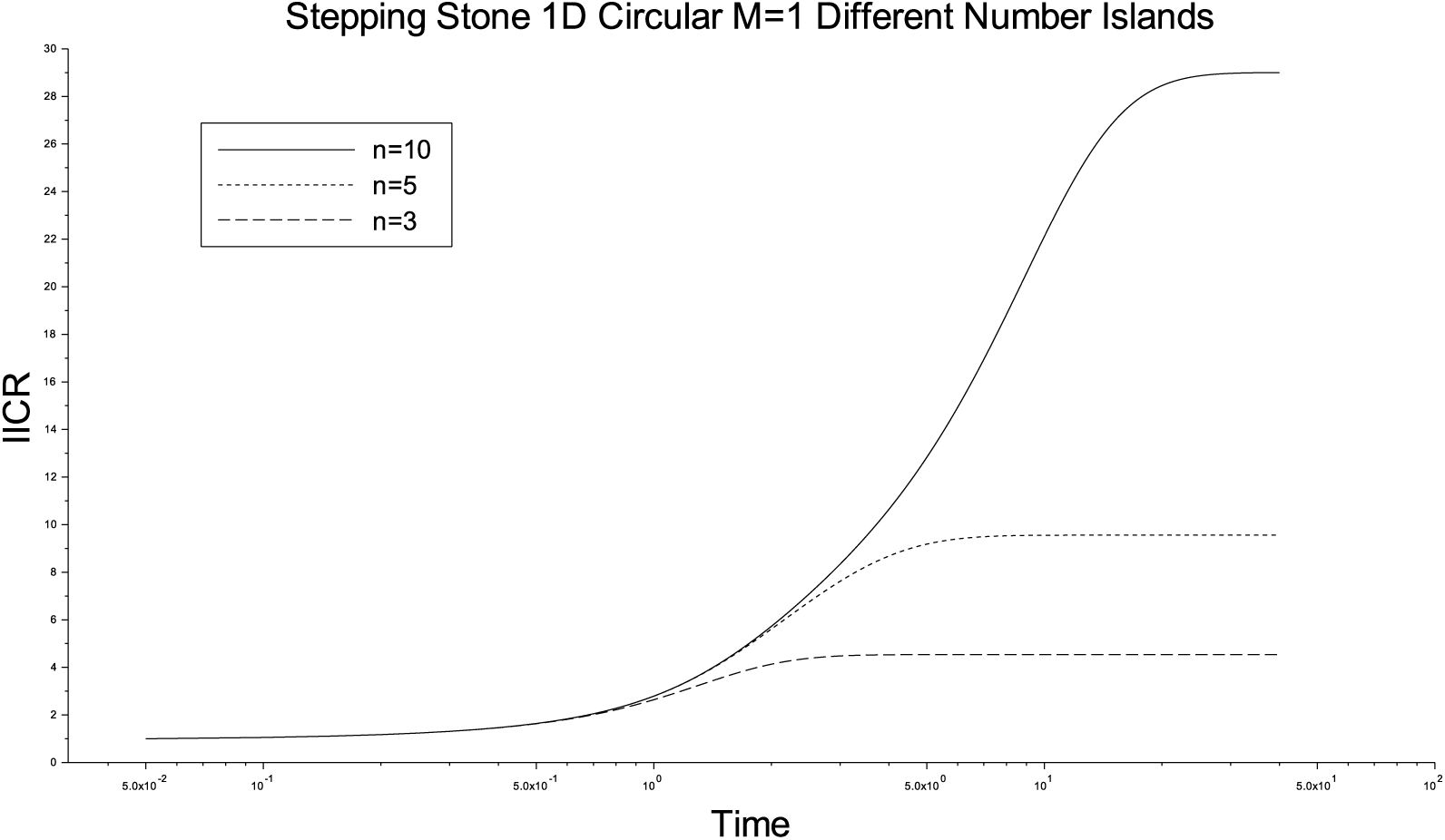
1D circular stepping stone, *M* = 1, different values of *n*

**Figure S5:**
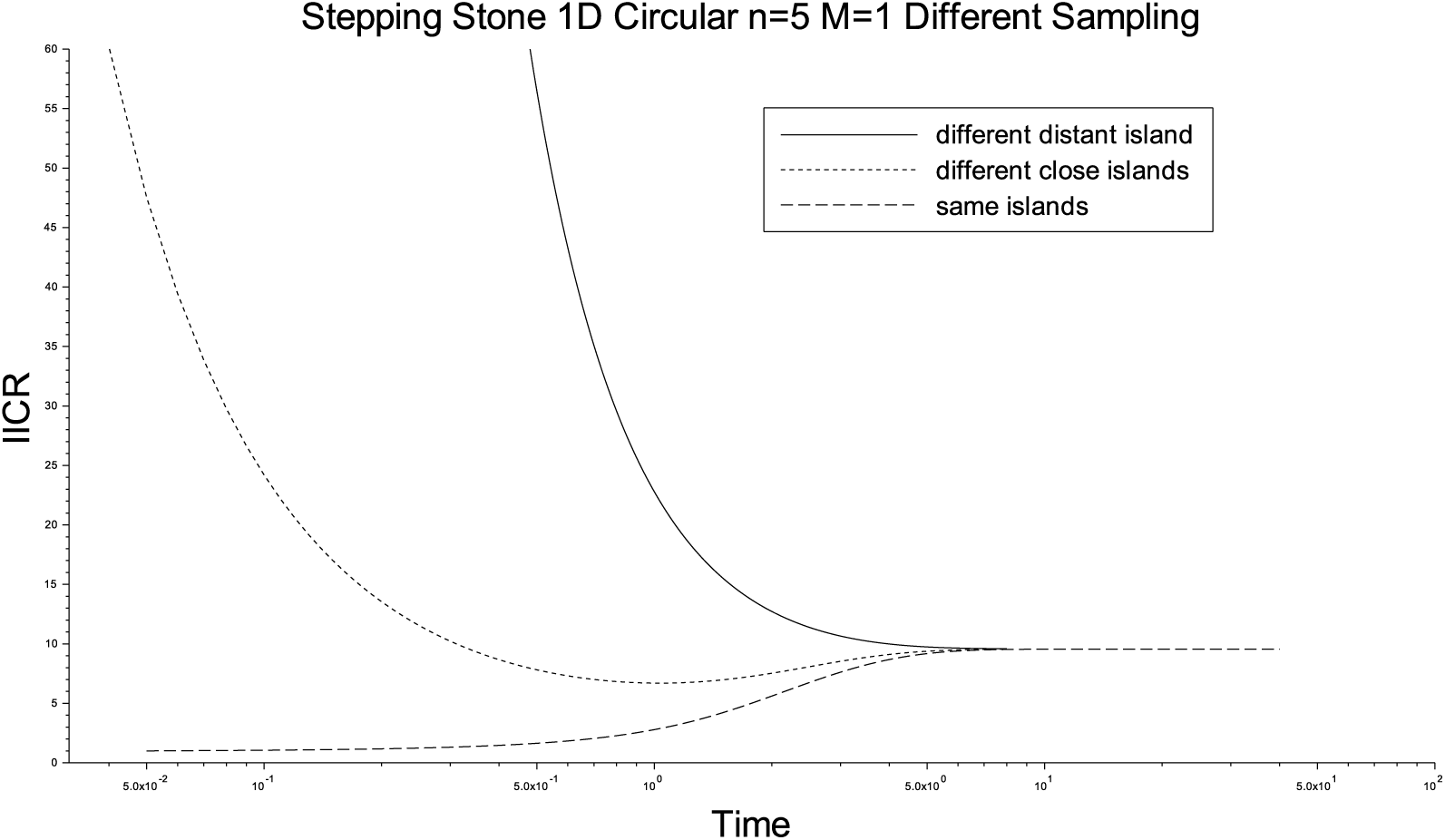
circular stepping stone, *n* = 5, *M* = 1, different sampling : two lineages in the same island, two lineages in nearby islands, two lineages in distant islands

**Figure S6:**
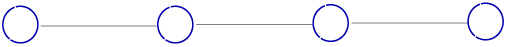
1D stepping stone with 4 islands

**Figure S7:**
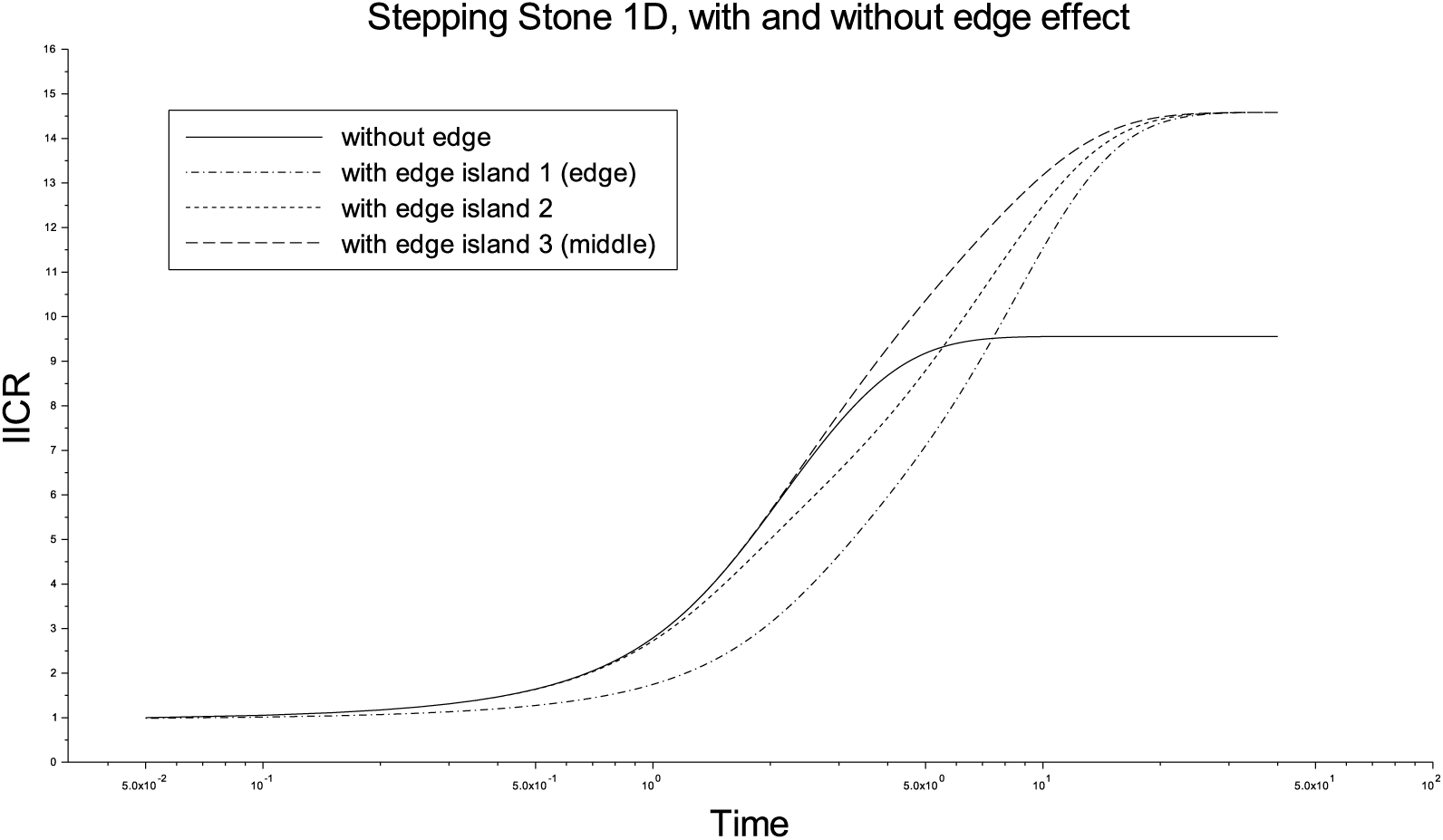
Comparison of two 1D Stepping Stone Models: with and without edge. Number of demes *n* = 5 and gene flow *M* = 1. Sampling two lineages in the same deme. When there is edge effect, we present the three ways to sample in the same island: extreme deme (number 1 or 5), demes right next to the extreme (2 or 4) and the middle one (number 3).

**Figure S8:**
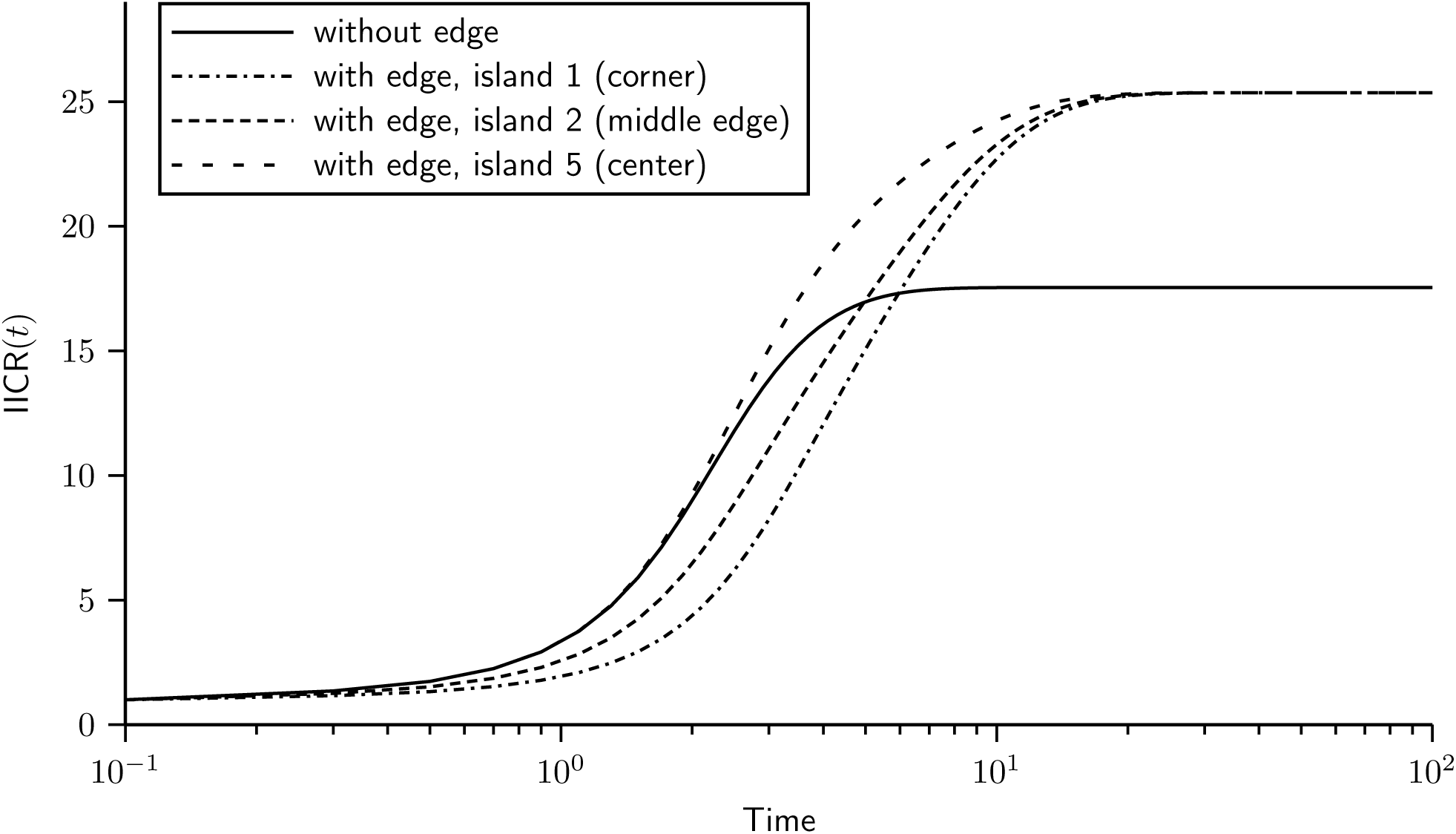
IICR plots for the 2D stepping stone model. Here we assumed a model with 3 × 3 = 9 islands and *M* = 1, with and without edge effect. In the model with edge effect, we plot the three ways to sample two lineages in the same island: in island 1, 3, 7 or 9 (corner), in island 2, 4, 6 or 8 (middle of the edge), and in island 5 (center of the lattice).

**Figure S9:**
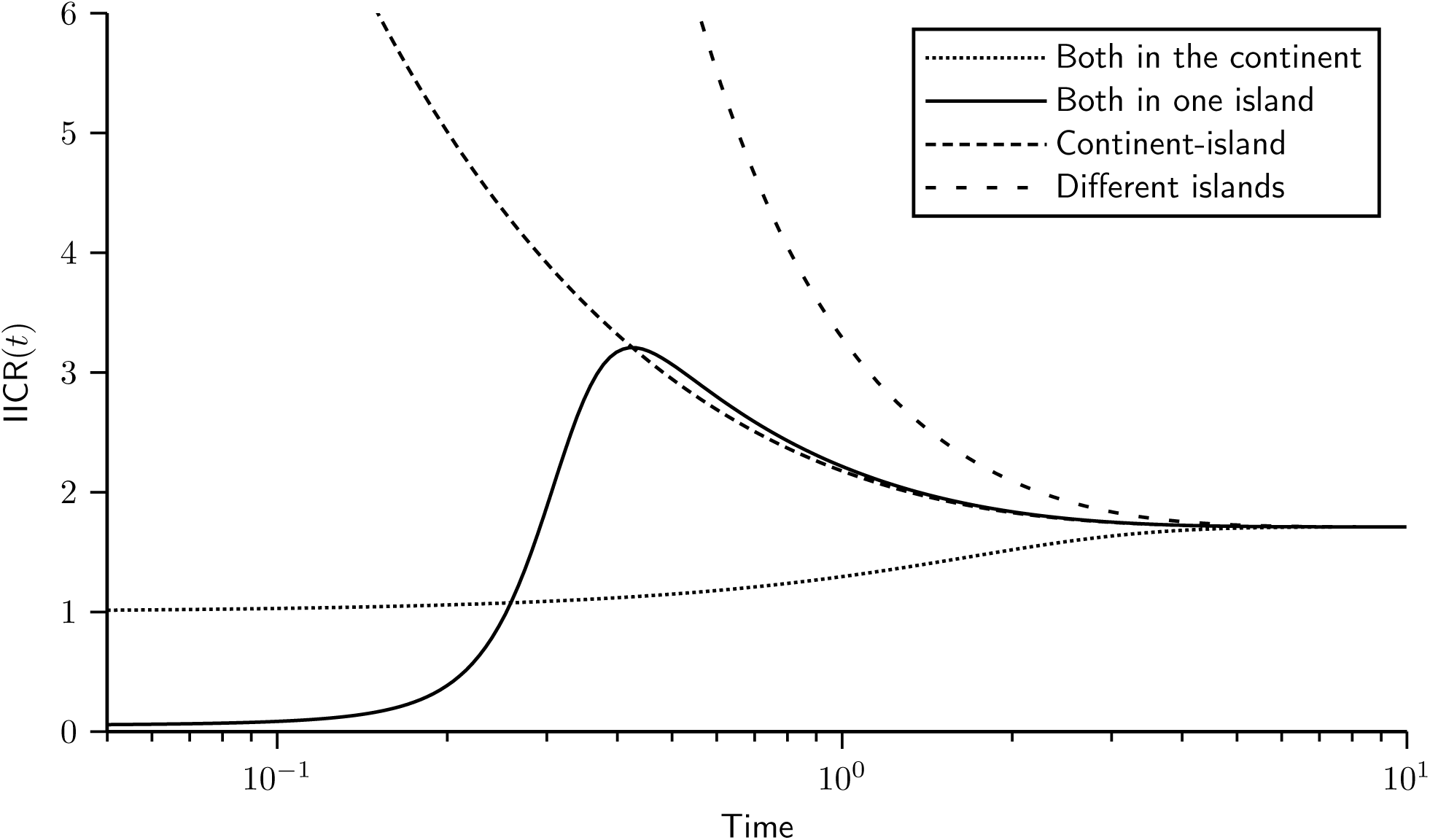
IICR for a continent-island model. We constructed the transition rate matrix for a model with *n* = 4, namely one continent and three same-sized islands. The sizes of the continent and of the islands were set to *c*_1_ = 1 and *c*_2_ = 0.05, respectively. In other words, the continent was 20 times larger than the islands. We set the migration rates to *M*_1_/2 = 0.05, *M*_2_/2 = 1 (note that once *M*_1_ is set, *M*_2_ is constrained to keep inward and outward migrant gene numbers equal, as required by equation 1). In this model there are only four types of IICR curves, two IICR_s_ and two IICR_d_. The first two correspond to the cases where we sample the two lineages either in the continent or in one of the islands. The IICR_d_ curves correspond to cases where one gene comes from the continent and the other from an island or when the two genes come from two different islands.

